# High-throughput living-cell protein crosslinking analysis uncovers the physiological relevance of forming the “inserted” state of the ATP synthase ε-subunit

**DOI:** 10.1101/749747

**Authors:** Yang Liu, Jiayu Yu, Mengyuan Wang, Qingfang Zeng, Xinmiao Fu, Zengyi Chang

**Affiliations:** State Key Laboratory of Protein and Plant Gene Research, School of Life Sciences, Center for Protein Science, Peking University, Beijing, 100871, China

## Abstract

ATP synthase, a highly conserved multi-subunit enzyme complex having a common stoichiometry of α_3_β_3_γδεab_2_c_8-15_, functions to supply ATP as the universal energy currency for cells. It comprises of the peripheral F_1_ sector (α_3_β_3_γδε) and the membrane-integrated F_o_ sector (ab_2_c_8-15_). *In vitro* structural analyses revealed that the C-terminal domain of the ε-subunit could adopt either an “inserted” or “non-inserted” state (with or without interacting with the α/β-subunits), with the former being viewed as inhibitory for the ATP hydrolysis activity of ATP synthase. Nevertheless, as common in current protein researches, the physiological relevance of such an “inserted” state for ATP synthase functioning is hardly known. To decipher this, designed an unnatural amino acid-mediated living-cell protein photocrosslinking analysis pipeline by developing the scarless genome-targeted site-directed mutagenesis and the high-throughput gel polyacrylamide gel electrophoresis (HT-PAGE) techniques. Employing this powerful approach, we systematically examined the interactions involving the C-terminal helix of the ε-subunit in cells living under a variety of experimental conditions. These studies enabled us to uncover that the “inserted” and “non-inserted” states of the ε-subunit exist as an equilibrium in cells cultured under common experimental conditions, shifting to the former upon the appearance of unfavorable conditions, acting as a low-gear state to strengthen the ATP synthesis function. Such a fine-tuning mechanism allows the ATP synthase to reversibly and instantly switch between two functional states. Further, the two powerful techniques that we developed here might be applied to many aspects of protein researches.

## Introduction

A constant energy supply, under conditions favorable or unfavorable, is vital for maintaining life. The ATP molecule, as the universal energy currency in cells, is commonly reproduced via substrate-level and/or oxidative phosphorylation (as well as photophosphorylation in photosynthetic organisms) of the ADP molecules. Oxidative phosphorylation (or photophosphorylation) occurs via the functioning of the highly conserved ATP synthase complex, which transforms the energy stored in the transmembrane proton gradient into that stored in the ATP molecule (Mitchell, 1961; Boyer, 1997; Walker, 2013). As the key player for oxidative phosphorylation, the ATP synthase is a multi-subunit enzyme complex that has a common stoichiometry of α_3_β_3_γδεab_2_c_8-15_. It consists of the peripheral F_1_ sector having the α_3_β_3_γδε subunits and the membrane-integrated F_o_ sector having the ab_2_c_8-15_ subunits (Boyer, 1997; Walker, 2013). Extensive studies have demonstrated that ATP synthesis, occurring at the F_1_ sector (with the active sites mainly contributed by the β-subunits), is driven by the down-hill proton movement across the F_o_ sector via a unique but not yet fully characterized (especially in living cells) rotational catalytic mechanism (Boyer, 1997; Walker, 2013).

Previous studies have demonstrated that both the isolated F_1_ sector and the holoenzyme of the ATP synthase exhibits ATP hydrolysis activity under *in vitro* conditions. The ATP synthase was therefore also called the F_1_F_o_-ATPase and may act in reverse to hydrolyze ATP for generating the transmembrane proton gradient in bacterial and other cells under particular conditions (Walker, 2013). These two opposite activities apparently would have to be regulated in way or another in adapting to different cellular conditions. In this regard, the ε-subunit has been described to play a role in suppressing the ATP hydrolysis activity of the ATP synthase, but largely based on *in vitro* observations (Laget and Smith, 1979; Sternweis and Smith, 1980; Nakanishi-Matsui et al., 2016).

Recent studies with X-ray crystallography, nuclear magnetic resonance (NMR) and cryoelectron microscopy revealed that the ε-subunit consists of two domains, the N-terminal domain that mainly exists as a ten-strand β-sandwich and the C-terminal domain as two α-helices (Wilkens and Capaldi, 1998; Cingolani and Duncan, 2011; Sobti et al., 2016). Moreover, the C-terminal domain of the ε-subunit has been observed to exist either as two extended α-helices, which would allow the C-terminal helix to interact with the α-, β-subunits in the F_1_ sector (Cingolani and Duncan, 2011; Sobti et al., 2016) could thus be designated as the “inserted” state (refer to **Fig. 1a**), or as a compact hairpin structure with the two α-helices lie side by side, which would not allow the C-terminal helix to interact with the α/β-subunits (Yagi et al., 2007; Sobti et al., 2019) and could thus be the “non-inserted” state.

**Figure 1.**
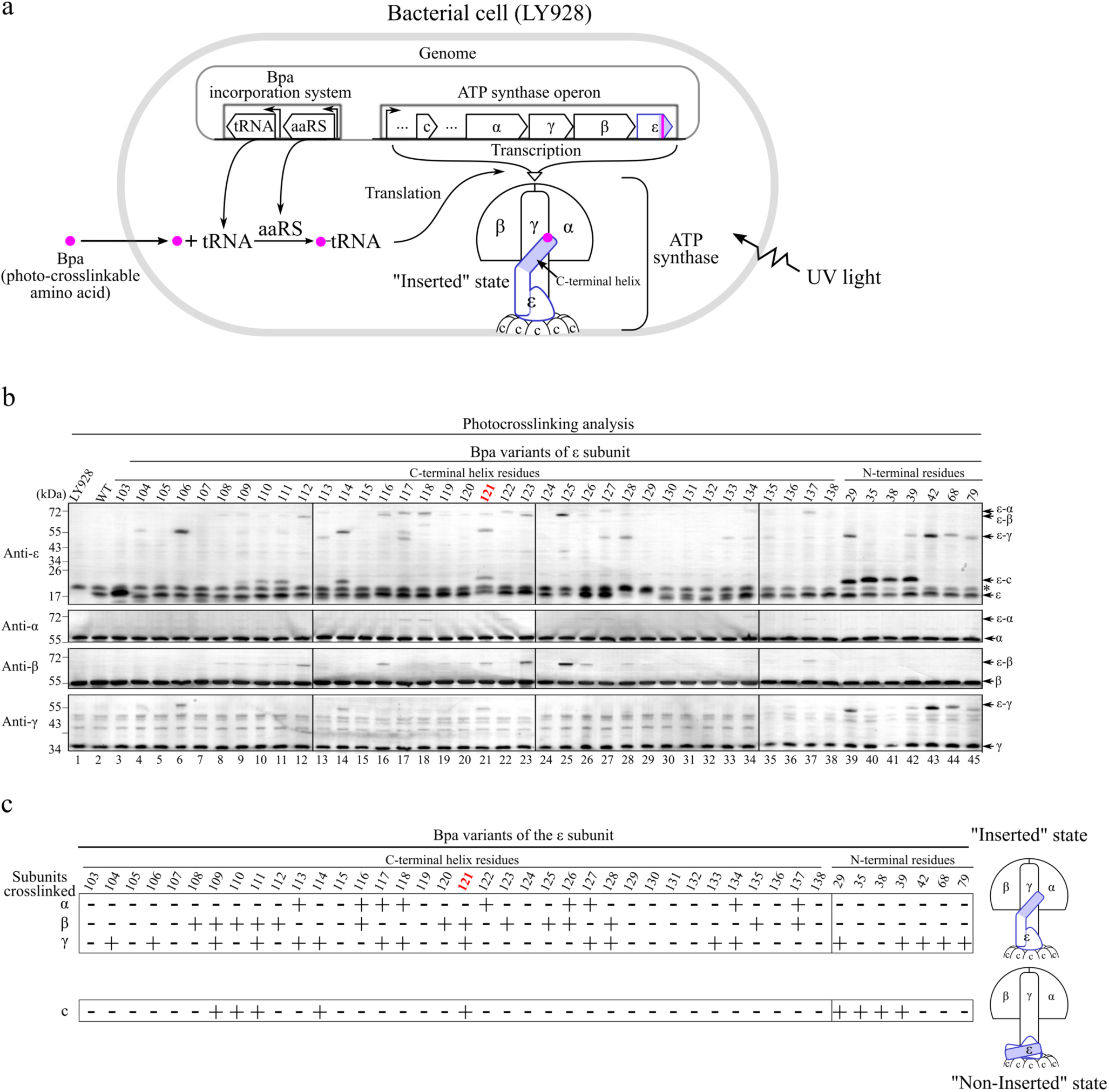
*In vivo* photocrosslinking analyses of Bpa variants as produced by using scarless genome-targeted site-directed mutagenesis reveal the formation of both the “inserted” and “non-inserted” states of the ε-subunit in bacterial cells overnight cultured in common LB rich medium. (**a**) Schematic diagram illustrating how a Bpa residue (pink dot) was introduced into the ε-subunit via transcription and translation after a selected codon was replaced by a TAG amber codon (pink bar) by employing the scarless genome-targeted site-directed mutagenesis technique (illustrated in more details in **Fig. S1a**). The LY928 bacterial strain utilized here was constructed by integrating the genes encoding the orthogonal amino-acyl tRNA synthetase (aaRS) and orthogonal tRNA, both needed for incorporating the unnatural amino acid Bpa, into its genomic DNA(Wang et al., 2016; Yu et al., 2019). An irradiation of the cells by UV light (for 30 sec to 10 min) would allow Bpa-mediated crosslinking to occur in living cells. (**b**) Blotting results for detecting the photocrosslinked products of the indicated Bpa variants of the ε-subunit (with multiple gels), probed with the alkaline phosphatase-conjugated streptavidin against the biotin linked to the Avi tag on the ε-subunit (Anti-ε), or antibodies against the α- (Anti-α), β- (Anti-β) or γ- (Anti-γ) subunit. The 7 Bpa variants produced at the N-terminal domain were analyzed here (lanes 39-45) for validating the feasibility of the scarless genome-based site-directed mutagenesis technique. Here the wild type (WT) cell (lane 2) was derived from the LY928 strain (lane 1) by having an Avi-tag connnected to the N-terminal end of the ε-subunit. Positions of the monomers of the α-, β-, γ-, ε- subunits, as well as crosslinked products between ε and other subunits (ε-α, ε-β, ε-γ, ε-c) are indicated on the right of the gels; positions of molecular weight markers are indicated on the left; * indicates a non-specific protein band detected with the alkaline phosphatase-conjugated streptavidin. (**c**) Summary of the crosslinking results for all the Bpa variants of the ε-subunit displayed in (**b**), indicating the formation of the “inserted” and the “non-inserted” states.

In one study, it was demonstrated that the extended “inserted” state of the ε-subunit could be converted to the compact “non-inserted” state in presence of a high concentration of ATP (Yagi et al., 2007; Sobti et al., 2019). Moreover, enzymatic activity analysis based on cysteine-mediated crosslinked forms suggested that the “inserted” state represents a form with which the ATP hydrolysis activity is inhibited (Tsunoda et al., 2001). Nonetheless, the truncation of the C-terminal helix of the ε-subunit was reported to exhibit a minimal impact on the growth or viability of the bacterial cells (Taniguchi et al., 2011). Because of these, a coherent explanation of the physiological context for such an inhibitory mechanism hitherto remains lacking (Walker, 2013; Nakanishi-Matsui et al., 2016). In other words, the physiologically meaningful condition under which such an “inserted” state of the ε-subunit is formed, as well as the biological role it plays, remain largely unknown. Generally speaking, correlating an interpretation of *in vitro* observations achieved using isolated proteins, which is often simple but sometimes out of the context, to a physiological context of living cells, which is often complex and dynamic in nature, remains a rather difficult thing in current explorations on proteins (and other biomolecules as well).

We might be able to decipher the physiological relevance of the “inserted’ state if we could directly and effectively characterize the interactions involving the C-terminal helix of the ε-subunit in cells living under a great variety of possible experimental conditions. In this regard, we decided to test our luck by performing *in vivo* protein photocrosslinking analysis on this C-terminal helix as mediated by the unnatural amino acid Bpa (p-benzoyl-L-phenylalanine), an approach that was initially developed by Schultz and colleagues (Chin et al., 2002), and has been extensively applied in our lab with a few technical modifications and improvements(Fu et al., 2019, 2013; Fu and Chang, 2019; Wang et al., 2016; Zhang et al., 2011). This approach recently allowed us to unveil how the FtsZ protein, a key cell division protein that has been extensively studied under *in vitro* conditions (Matsui et al., 2012; Li et al., 2013), assembles into the dynamic Z-ring structure via novel lateral interactions of its protofilaments in actively dividing living bacterial cells (Guan et al., 2018) but is sequestered in an reversible subcellular structure that we named as the regrowth-delay bodies in non-dividing/non-growing cells (Yu et al., 2019).

For this study, we developed two new techniques which enabled us to effectively capture the dynamic protein-protein interactions involving the C-terminal helix of the ε-subunit in cells living under a great variety of experimental conditions in high spatial (at the level of amino acid residues) and temporal (at the level of minutes) resolutions. In particular, the scarless genome-targeted site-directed mutagenesis technique allowed us to introduce the Bpa residue into the ε subunit by modifying its endogenous gene in the genomic DNA (rather than by modifying an exogenous gene carried by a recombinant expression vector, as conventionally done). It follows that the Bpa variants were expressed with the timing and quantity of production to be identical to that of the wild type ε-subunit. More importantly, the high-throughput polyacrylamide gel electrophoresis (HT-PAGE) technique enabled us to perform Western blotting analysis of up to 384 protein samples per single gel. It follows that the photocrosslinked products of all the Bpa variants of the ε-subunit, reflecting the interaction participated by the replaced residues, could be examined in a semi-quantitative fashion.

With these two new techniques, we individually introduced the unnatural amino acid Bpa at all the 36 residue positions in the C-terminal helix of the ε-subunit and then performed high-throughput analysis on the crosslinked products formed by these variant proteins that were expressed in the authentic genome context and in living cells placed or cultured under a great variety of experimental conditions. These analyses allowed us to unveil, strikingly, that the ε-subunit exists as the “inserted” state under such common experimental conditions as when the cells are cultured in the LB rich medium. Further studies enabled us to reveal that the ε-subunit exists as the “inserted” and “non-inserted” states in an equilibrium and can effectively switch between them depending on the environmental conditions, with the “inserted” state to be predominant when the cells are facing such unfavorable conditions as in the moderate presence of a natural or non-natural uncoupler that weakens the transmembrane proton gradient. By effectively identifying a unique mutant (I125K) of the ε-subunit that was severely defective in forming the “inserted” state, we demonstrated that the “inserted” state apparently functions to strengthen the ATP synthase, which in turn provides the cell a moderate but significant advantage for growing under unfavorable conditions. The switch between these two quaternary structure states seems to act for fine-tuning the function of the ATP synthase via a reversible and instant manner, which is somehow analogous to gear-shifting in an automobile: allowing the enzyme complex to switch to the “inserted” state, representing a low-gear strengthened state when the cells are living under unfavorable conditions but to the “non-inserted” state, representing a high-gear non-strengthened state under favorable ones. It is possible that such a fine-tuning mechanism is employed by many enzymes or enzyme complexes, which would allow cells to rapidly adapt to changing living conditions. In addition, the two powerful techniques that we developed here possess potential applications not only in living-cell protein photocrosslinking analysis but also in many other aspects of protein researches.

## Results

### Protein photocrosslinking coupled with scarless genome-targeted site-directed mutagenesis reveal that the “inserted” state of the ATP synthase ε-subunit could be detected in living bacterial cells overnight cultured in common LB rich medium

In an attempt to find out whether the “inserted” state of the ε-subunit of the ATP synthase is actually formed in living cells, we focused on the interactions involving the C-terminal helix (i.e., the α-helix at the C-terminal end) of the ε-subunit. For this purpose, we made use of the photoactivatable unnatural amino acid p-benzoyl-L-phenylalanine (Bpa), which after being genetically incorporated into a target protein and upon UV irradiation would be able to effectively mediate covalent photocrosslinking with the interacting proteins in cells living under any particular experimental conditions (Chin et al., 2002; Zhang et al., 2011; Fu et al., 2013; Wang et al., 2016; Yu et al., 2019; Fu and Chang, 2019).

Moreover, to maintain the strict stoichiometry of all the subunits of the ATP synthase complex (which would be highly desirable in studying any multi-subunit protein), we developed a scarless genome-targeted site-directed mutagenesis technique to produce the Bpa variants of the ε-subunit by modifying its endogenous gene in the genomic DNA, as schematically illustrated in **Fig. 1a**. Briefly, a selected codon in the ε encoding gene was replaced by a TAG amber codon via a procedure involving two rounds of homologous recombination (illustrated in more details in **Fig. S1a**). This would produce a specific Bpa variant of the ε-subunit (or any other protein) under the authentic genomic context such that its timing and quantity of production are completely identical to that of the wild type ε-subunit. It should be pointed out that all the Bpa variants of the ε-subunit described here were demonstrated to be functional, judging from the fact that they all allowed the modified cells to effectively grow in the succinate-containing M9 minimal medium (**Fig. S1b**), in which the functioning of the ATP synthase was indispensable for the cells to grow. In addition, for an effective introduction of Bpa into target proteins, here we made use of the *E. coli* bacterial strain LY928 that we generated by inserting the optimized DNA sequences encoding the orthogonal Bpa-tRNA synthetase and the orthogonal tRNA^Bpa^ at a specifically selected site on the genomic DNA causing hardly any disruption on cellular activities (Wang et al., 2016; Yu et al., 2019).

For the purpose of effective detection, we also modified the genomic ε-encoding gene such that an Avi-tag (Gräslund et al., 2017) was added to the N-terminal end of the ε-subunit. This Avi-ε form was proven to allow the modified cells to grow in the succinate-containing M9 minimal medium as effectively as the unmodified LY928 cells (**Fig. S1b**), indicating that the Avi-ε form was effectively assembled into a fully functional ATP synthase complex. In light of this and for the convenience of description, we hereafter designate this Avi-ε form as the wild type form of the ε-subunit, such that all the Bpa variants of the ε-subunit we generated in this study would differ from this wild type form by merely having a selected amino acid residue replaced by a Bpa residue.

We first tested the feasibility of this genome-targeted site-directed mutagenesis technique by individually introducing Bpa to seven residue positions in the N-terminal domain of the ε-subunit. This is because of the fact that the structure as well as the interactions involved for this domain, unlike that for the C-terminal domain, have been known to be relatively less dynamic (rather than highly dynamic), based on the information provided by *in vitro* structural analyses. In particular, four residues (at positions 29, 35, 38 and 39) were reported to interact with the c-subunit while the other three (at positions 42, 68 and 79) interacting with the γ-subunit of the ATP synthase (Cingolani and Duncan, 2011; Sobti et al., 2016).

Results of blotting analysis, as presented in **Fig. 1b**, demonstrated that all the ε variants having a Bpa introduced at residue position 29, 35, 38 and 39 (lanes 39-42) indeed formed crosslinked products (thus interacting) with the c-subunit, while those having a Bpa introduced at position 42, 68 and 79 formed crosslinked products with the γ-subunit (lanes 43-45) in living bacterial cells cultured overnight in common LB rich medium. Additionally, we observed that the Bpa introduced at residue position 29 or 39 was able to mediate an interaction not only with the *c* but also with the γ-subunit (lanes 39 and 42, respectively, in **Fig. 1b**). This was in agreement with the fact that these two residues are located in the ε-subunit close to a cross point of the γ- and c*-*subunits in the ATP synthase (Sobti et al., 2016). Collectively, these observations demonstrated that each of these substituted Bpa residues was correctly introduced via the genome-targeted site-directed mutagenesis. This validated the effectiveness of our strategy (as shown in in **Fig. 1a**) for characterizing and profiling the quaternary structures of the C-terminal helix of the ε-subunit in living cells, with a structural resolution at the level of individual amino acid residues.

We then applied this scarless genome-targeted site-directed mutagenesis technique to individually introduce Bpa at all the 36 residues (103-138) of the C-terminal helix of the ε-subunit and subsequently analyzed their photocrosslinked products formed in living bacterial cells. With no idea on the experimental condition under which the C-terminal helix might form the “inserted” state in living cells, we did an initial random search by performing photocrosslinking analysis after these 36 variant cells were cultured overnight in common LB rich medium. Strikingly, we observed an effective formation of the “inserted” state of the ε-subunit even under such a common experimental condition, as reflected by the detection of crosslinked products between many of these Bpa variants (e.g., when Bpa was placed at residue position 121 or 125) of the ε-subunit and the α-, β- or γ-subunit (the top panel in **Fig. 1b**, also summarized in **Fig. 1c**). The nature of these crosslinked products, other than being roughly judged on the basis of the expected combined molecular masses of the two crosslinked subunits (i.e., ε-α, ε-β or ε-γ), were also verified by simultaneously probing the parallel gels with antibodies against the α-, β- or γ-subunit (**Fig. 1b**, the bottom three panels).

Remarkably, we meanwhile observed that multiple of these Bpa variants of the ε-subunit (e.g., when Bpa was placed at residue positions 109-111,114, and 121) were also able to form crosslinked products with the c*-*subunit (**Figs. 1b** and **1c**). We view these results, though with hardly any hint from previous *in vitro* structure studies, as indicating the simultaneous presence of the “non-inserted” state of the ε-subunit in these cells. Here, the band representing the crosslinked product between ε- and c*-*subunits (lanes 9, 10, 11, 14 and 21 in **Fig. 1b**) was identified by its mobilization to the same position as those formed between the ε- and c-subunits when Bpa was introduced at residue position 29, 35, 38 or 39 (lanes 39, 40, 41 and 42 in **Fig. 1b**).

Interestingly, we also noticed that Bpa residues introduced at some of these positions (e.g., 114 and 121) mediated interactions not only with the α/β/γ-subunits but also with the c-subunit. This apparently implies that residues at these positions mediate the formation of both the “inserted” and the “non-inserted” states of the ε-subunit. In view of the fact that the α- and β-subunits are at least ~40 Å away from the *c* subunit, one particular residue in the C-terminal helix of a ε-subunit would be impossible to simultaneously interact with both the α/β-and c-subunits in one single ATP synthase complex. One plausible explanation of these results is that, under this particular experimental condition, the “inserted” state might exist in some cells while the “non-inserted” exist in other cells, or alternatively, the two states might simultaneously exist in different ATP synthase complexes of one single cell.

These better-than-expected observations stimulated us to further explore the exact physiological contexts in which the “inserted” state of the ε-subunit might become the predominant in cells. Further, whether the “inserted” state is able to be converted back to the “non-inserted” state, and vice versa, under certain experimental conditions? We reasoned that, the most practical way to address these issues was to treat the cells under a variety of experimental conditions and meanwhile to monitor the interaction patterns of the C-terminal helix of the ε-subunit by examining the photocrosslinked products formed by all the 36 Bpa variants. In theory, this might provide a powerful way of self-validations, since a particular interaction pattern involving the C-terminal helix should be simultaneously reflected by many of these Bpa variants. However, for the blotting results of all the Bpa variants to be compared in a precise manner, all (of hundreds) the protein samples would be ideally resolved via exactly the same experimental (electrophoresis and blotting) operation, i.e., on one single gel. To the best of our knowledge, such a high-throughput gel electrophoresis and blotting technique remains unavailable and we decided to develop one.

### Development of a high-throughput polyacrylamide gel electrophoresis (HT-PAGE) technique that allows up to 384 protein samples to be effectively analyzed on one single gel

We started the efforts of developing this high-throughput technique with the rough idea of converting the current vertical form of the sodium dodecyl sulfate polyacrylamide gel electrophoresis (SDS-PAGE) to a submerged horizontal one, similar to how the agarose gel electrophoresis has been applied to DNA analysis. We reasoned that such a horizontal gel system, unlike a vertical one, might allow us to increase the number of protein samples analyzed per gel by adding more rows of sample-loading wells. The most challenging issue in developing such a technique would be to achieve a resolution high enough to distinguish protein bands having close molecular masses, for instance, the ε-subunit and the ε-c crosslinked product (as shown in **Fig. 1b**).

In retrospect, we developed this submerged horizontal high-throughput polyacrylamide gel electrophoresis (HT-PAGE) technique via roughly four trial-and-error stages of designs and improvements, sometimes without understanding why certain experimental operations worked while others failed. This was somehow analogous to how Koichi Tanaka developed the matrix-assisted laser desorption/ionization time-of-flight mass spectrometry for protein analysis(Tanaka, 2003). As shown in **Fig. 2** (schematic illustration on the left, actual blotting results on the right), the scaling up of the system along the four stages was accomplished by a stepwise increase in the number of sample lanes per gel from 15 (at stage 1) to 30 (at stage 2) to 60 (at stage 3) and eventually to 384 (at stage 4). This increase in sample lanes was made possible mainly as a result of a proportional decrease of the actual size for each sample lane, from 3 × 20 mm to 3 × 10 mm to 1 × 10 mm and eventually to 1 × 8 mm (**Fig. 2**). As demonstrated here, the resolution of the blotting results in the 384-sample per-gel system was largely comparable with that of the 15-sample per-gel system, both allowing all the major crosslinked products of the ε-subunit to be clearly distinguishable.

**Figure 2.**
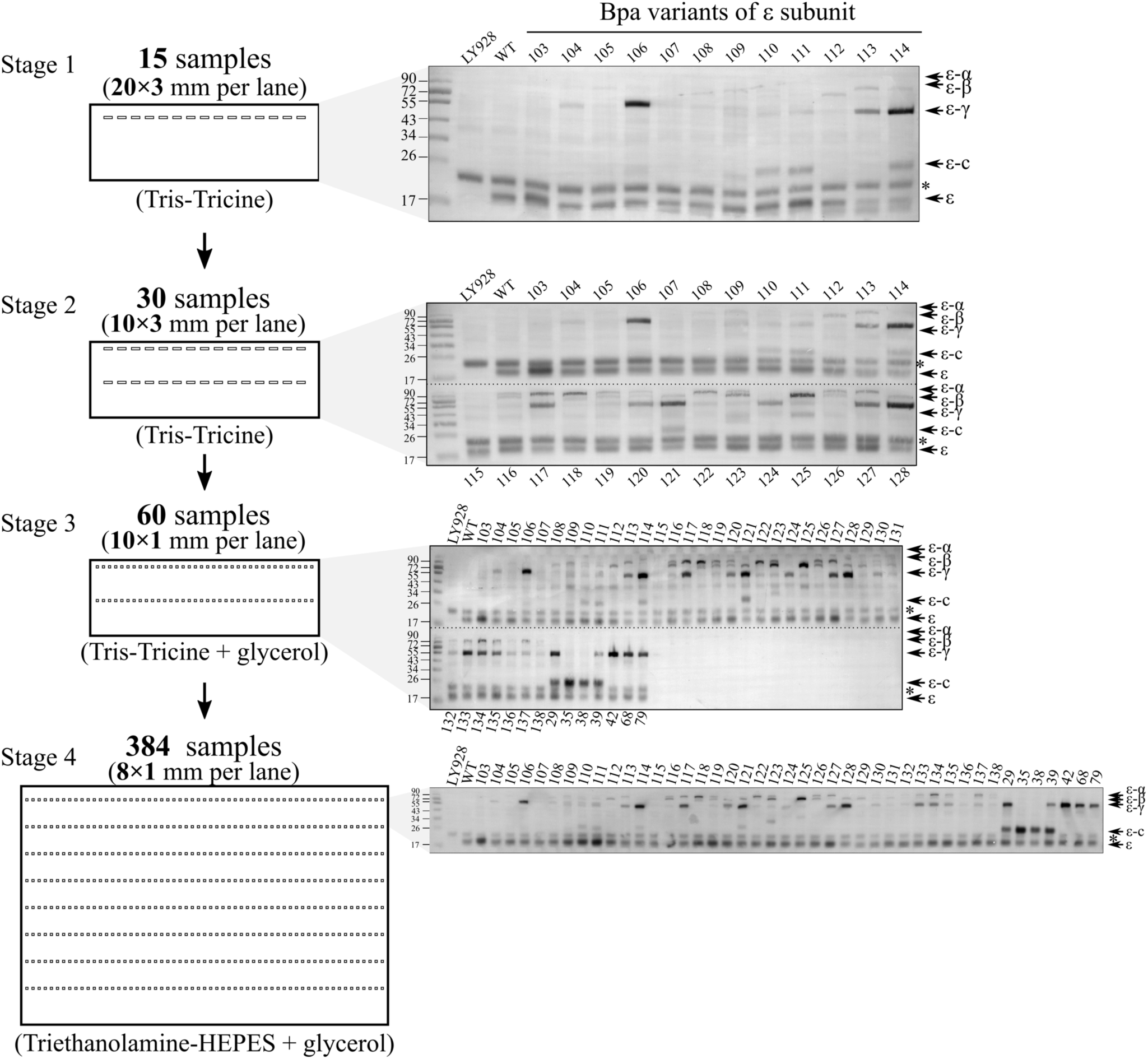
Development of the high-throughput SDS polyacrylamide gel electrophoresis (HT-PAGE) technique that enables one to effectively analyze up to 384 protein samples on one single gel. Shown on the left is a schematic illustration of the four stages we went through in developing HT-PAGE and on the right displays the corresponding actual blotting results obtained at each stage. For each stage of development (on the left), the following parameters are indicated: the number of protein samples analyzable per gel, the size of each gel lane, and the buffer that worked; again, * indicates a non-specific band. The crosslinked products analyzed at each stage were taken from those analyzed in Fig. 1b (only part of them for stages 1 and 2, all of them for stages 3 and 4).

In sum, the successful development of the HT-PAGE, a detail protocol of which is described in detail in the Materials and Methods, was mainly resulted from the following designs and explorations. First, finding a proper buffer for preparing and running the gel, though extremely time consuming and rather challenging, was proven to be one critical aspect for this success. After testing many different running buffers, we found that although Tris-tricine worked effectively for Stages 1 through 3, only TEA (triethanolamine)-HEPES (4-(2-hydroxyethyl)-1-piperazineethanesulfonic acid) enabled us to achieve a high resolution at Stage 4, i.e., for the final HT-PAGE. Second, for Stages 3 and 4, glycerol had to be added into the buffer to increase the viscosity of the samples, thus preventing the occurrence of sample diffusion due to the small amount loaded as well as the longer time taken for loading all the hundreds of protein samples. Third, a ratio of the acrylamide cross-linking agent (usually N, N’-methylenebisacrylamide) /total acrylamide of 6% (w/w) was also found to be critical for preparing an effective polyacrylamide gel. Fourth, the distance interval between each two neighboring row of sample wells in the 384-lane gel had to be 8 mm for the samples to be loaded using a commercially available eight-channel pipet (because the distance between each two neighboring channels is 8 mm). Collectively, the HT-PAGE technique that we developed here enables one to analyze up to 384 protein samples (of any nature) in one single gel such that the levels of any specific protein band could be effectively compared in a semi-quantitative fashion across all the lanes.

### The “inserted” state of the ε-subunit are most predominant in the overnight cultured LB rich medium and in the CCCP-containing M9 minimal medium but becomes hardly detectable in fresh LB rich medium and glucose-containing M9 minimal medium

We next attempted to identify the experimental conditions under which the “inserted” state of the ε-subunit is the most predominant. For this purpose, we subjected all the 36 variant cells, collected from overnight cultured LB rich medium, to a short duration of treatment (~10 min) in a great variety of solutions and monitored the level of the “inserted” (and “non-inserted”) state via the relative density of the crosslinked ε-α, and/or ε-β, and/or ε-γ product bands across all the 36 Bpa variants (the whole operation pipeline is schematically illustrated in **Fig. 3**).

**Figure 3.**
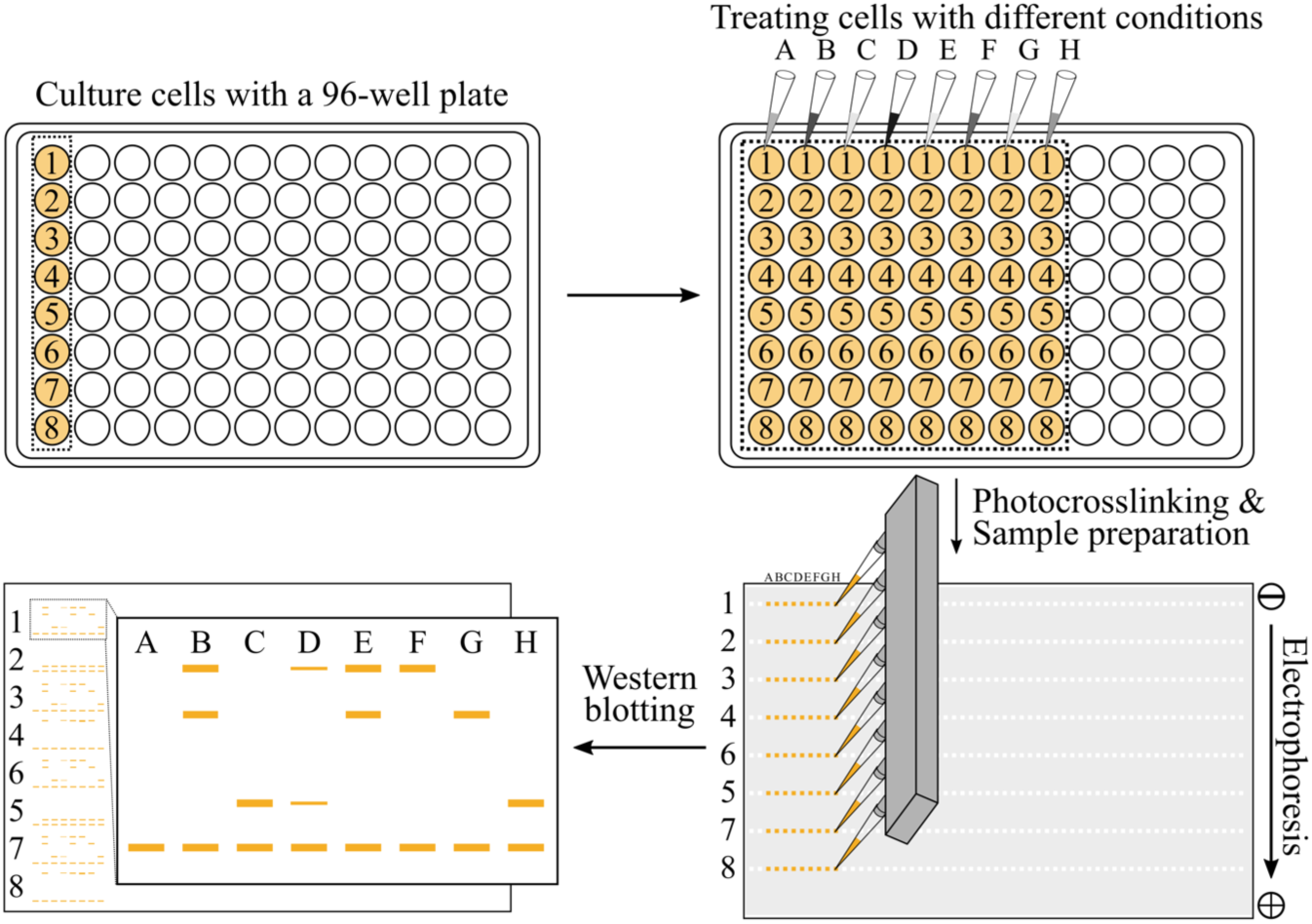
Schematic diagram illustrating how a large number of different cell samples could be cultured and treated in parallel under different experimental conditions and photocrosslinked before subjecting to HT-PAGE analysis. The 96-well culture plates were utilized for growing and treating the cell samples (arbitrarily shown here are eight different types of cells, each being treated under eight different conditions) and eight-channel pipet were used for effectively loading the protein samples onto the 384-lane gel. The Western blotting analysis was performed according to commonly available protocols without any modifications.

The blotting results for detecting the photocrosslinked products of the Bpa variants as examined with one single gel are displayed in **Fig. 4a**, which should be viewed as representing those derived from treatments with only a few selected solutions out of the many that we explored during this study. Here, the crosslinking products of each overnight-cultured Bpa variant cell were also analyzed (the *o^+^* lanes; though being identical in nature to those shown in **Fig. 1b**, the cells were now cultured using the 96-deep-well plate, rather than in test tubes as done above) on this high-throughput gel. For the sake of clarity, we will describe these results according to the rough logic we applied in designing these treatments. First, we simply treated the overnight cultured cells in fresh LB rich medium for ~10 min. Strikingly, we observed a significant reduction in the level of the “inserted” state upon such a simple solution change, as reflected by a significant reduction in the density of the ε-α and/or ε-β and/or ε-γ bands. This is evident by comparing the densities of the *f* and *o^+^* lanes for the 121-, 125-, 137- or any other Bpa variants in **Fig. 4a**, which meanwhile showed the power of our HT-PAGE technique. These interesting results strongly suggested that not only the formation of the “inserted” state is able to respond to changing environmental conditions but to respond in an instant manner. Moreover, it also indicated that certain component(s) present in the overnight cultured LB rich medium but not in the fresh LB rich medium might be critical for the formation of the “inserted” state of the ε-subunit. Nevertheless, we could not rule out the possibility that certain component(s) present in the fresh LB medium but not in the spent medium promoted the elimination of the “inserted” state (or the formation of the “non-inserted” state). This remarkable observation inspired us to further identify the more exact physiological conditions under which the “inserted” state of the ε-subunit would be formed (or more precisely, being present in a predominant manner).

**Figure 4.**
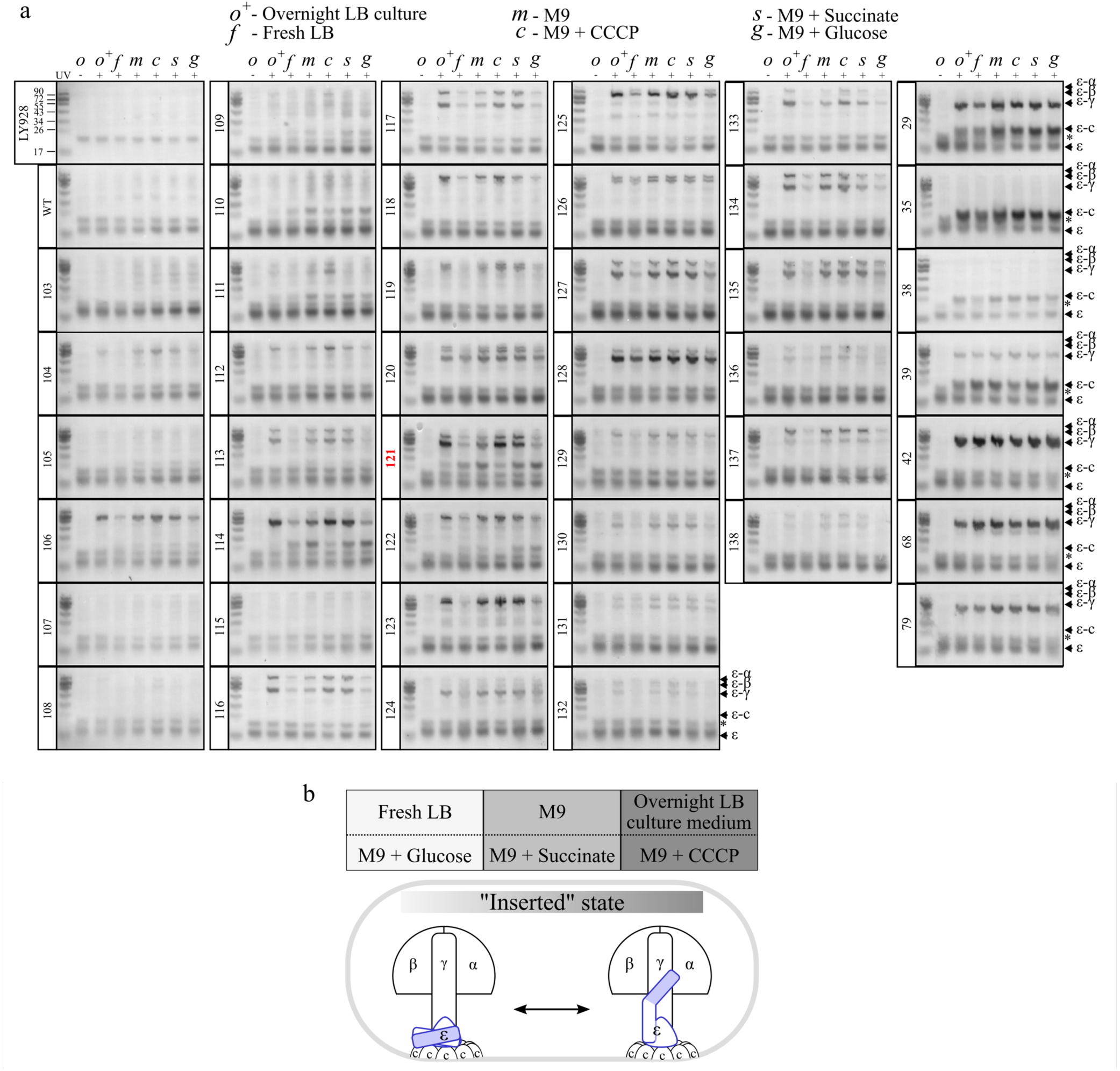
The “inserted” state of the ε-subunit is predominant in cells overnight cultured in LB rich medium or in cells treated in CCCP-containing M9 minimal medium but is bearly detectable when the cells are treated in fresh LB rich medium or glucose-containing M9 minimal medium. (**a**). Blotting results for detecting the photocrosslinked products of the 36 variants of the C-terminal helix (Bpa at residues 103-138) and the 7 variants of the N-terminal domain (Bpa at residues 29, 35, 38, 39, 42, 68, or 79) of the ε-subunit, after each type of the cell was overnight cultured in the LB rich medium and then treated in the indicated solutions for about 10 minutes, probed with alkaline phosphatase-conjugated streptavidin against the ε-subunit. The LY928 and wild type (Avi-ε LY928) cells were similarly treated and analyzed here as negative controls. Positions of the ε monomer or crosslinked ε products are indicated on the right of the gel; again, * indicates a non-specific band. (**b**). Summary of the relative levels of the “inserted” and “non-inserted” states of the C-terminal helix of the ε-subunit that were detected under the indicated experimental conditions, as indicated by the crosslinking data displayed in (**a**).

Because of the complex and undefined nature of the LB rich medium, we then decided to perform further studies with such solutions as the M9 minimal media, whose composition is much simpler and can be clearly defined. In particular, we then treated each of these 36 Bpa variant cells in M9 minimal medium (containing 47.7 mM Na_2_HPO_4_, 22.0 mM KH_2_PO_4_, 8.55 mM NaCl, 18.7 mM NH_4_Cl) alone, or in the M9 minimal medium containing 10 μM CCCP (carbonyl cyanide 3-chlorophenylhydrazone at a concentration of, a commonly used uncoupler that might immediately disrupt the transmembrane proton gradient of the cells), or in the M9 minimal medium containing 0.2% (w/v) succinate (which would allow the cells to undergo oxidative phosphorylation via respiration, i.e. to make the ATP molecules via the ATP synthase), or in the M9 minimal medium containing 0.2% (w/v) glucose (which might share certain feature as the fresh LB rich medium in allowing cells to initially produce ATP via substrate-level phosphorylation, before switching to the ATP synthase-participated oxidative phosphorylation).

The blotting results for detecting the crosslinked products that were immediately formed after the cells were treated in these various M9 minimal media (**Figs. 4a** and **4b**) revealed the following. First, when the cells were treated in M9 minimal medium with the addition of glucose (the *g* lanes), the level of the “inserted” state was very similar to that detected when the cells were treated in the fresh LB rich medium (the *f* lanes), i.e., being significantly lower than that detected in the overnight cultured LB rich medium. Second, when the cells were treated in the M9 minimal medium alone (the *m* lanes) or with the addition of succinate (the *s* lanes), the level of the “inserted” state was only slightly reduced but remained high comparing with that detected in cells derived from the overnight cultured LB rich medium (compare with the *o*^+^ lanes). Third, when the cells were treated with the CCCP-containing M9 minimal medium (the *c* lanes), interestingly, the level of the “inserted” state was largely comparable with that detected in the overnight cultured LB rich medium (the *o^+^* lanes). Collectively, these observations apparently demonstrated that certain component(s) is(are) present in the unfavorable overnight cultured LB rich medium, which played a role similar to CCCP (at 10 μM), both promoted a predominant formation of the “inserted” state for the ε-subunit in the bacterial cells.

Notably, during all these treatments (including those not shown in **Fig. 4a**), we never observed a complete absence of the “non-inserted” state (as represented by the formation of the ε-c crosslinked product). Similar to that of the “inserted” state, its level was also variable, being low in the overnight culture media and in CCCP-containing M9 minimal medium but high in the M9 minimal medium alone, or with the addition of either succinate or glucose. These observations once again supported our conclusion that the “inserted” and “non-inserted” states of the ε-subunit somehow exist as an equilibrium in the ATP synthase complexes. In addition to the 36 Bpa variants of the C-terminal helix of the ε-subunit, as a kind of control, we also analyzed the 7 Bpa variants of its N-terminal region, with the cells being treated with the same solutions. In contrast to the variable interactions involving the C-terminal helix, the interaction involving the N-terminal domain of the ε-subunit was largely unaltered upon treating with all these solutions (the right column in **Fig. 4a**). These results demonstrated that only the C-terminal domain of the ε-subunit undergoes a dynamic interaction with other subunits of the ATP synthase.

Collectively, as summarized in **Fig. 4b**, these *in vivo* crosslinking results revealed that the “inserted” state of C-terminal helix of the ε-subunit was predominant in cells present in the spent LB rich medium or the CCCP-containing M9 minimal medium, both being unfavorable conditions for the cells. However, it became non-predominant in cells placed in the fresh LB rich medium or glucose-containing M9 minimal medium, both being favorable conditions for the cells. Moreover, the “inserted” state seems to be able to convert to the “non-inserted” state in an instant fashion, in responding to a change to favorable environmental conditions.

### The “non-inserted” of the ATP synthase ε-subunit is automatically switch to the “inserted” state in bacterial cells upon culturing for a certain duration in fresh LB rich medium or in glucose-containing M9 minimal medium

Our observations described above demonstrated that the “inserted” state of the ε subunit as present in cells overnight cultured in LB rich medium could be immediately converted to the “non-inserted” state upon resuspension in fresh LB rich medium (**Fig. 4a**). It follows that there has to be a time-dependent conversion of the “non-inserted” state back to the “inserted” state for the cells during their culturing in the fresh LB rich medium. To verify this, we selected seven representative Bpa variant cells (with Bpa introduced at residue position 106, 110, 114, 121, 127, 133 or 137) of the C-terminal helix of the ε subunit and examined the photocrosslinked products formed in them after culturing them in fresh LB rich medium at 37°C for various durations.

The blotting results for detecting the photocrosslinked products, as also performed via HT-PAGE analysis, demonstrated the following. First, the level of the “inserted” state (reflected by the density of the ε- α and/or ε- β, and/or ε- γ crosslinked bands) was significantly increased after the cells were cultured in fresh LB rich medium for a short period of time, becoming a predominant form approximately at the 30 minutes time point (see the “LB” panels in **Fig. 5a**). Second, this time-dependent increase of the “inserted” state of the ε-subunit was similarly observed when the cells were cultured in the glucose-containing M9 minimal medium, only that it took about one hour for the “inserted” state to became a predominant (the “M9 + glucose” panels in **Fig. 5a**). Third, when the cells were cultured in the succinate-containing M9 minimal medium, the “inserted” state was significantly detected even at the 0 time point and became predominant at about the 1.5 h time point (the “M9 + succinate” panels in **Fig. 5a**).

**Figure 5.**
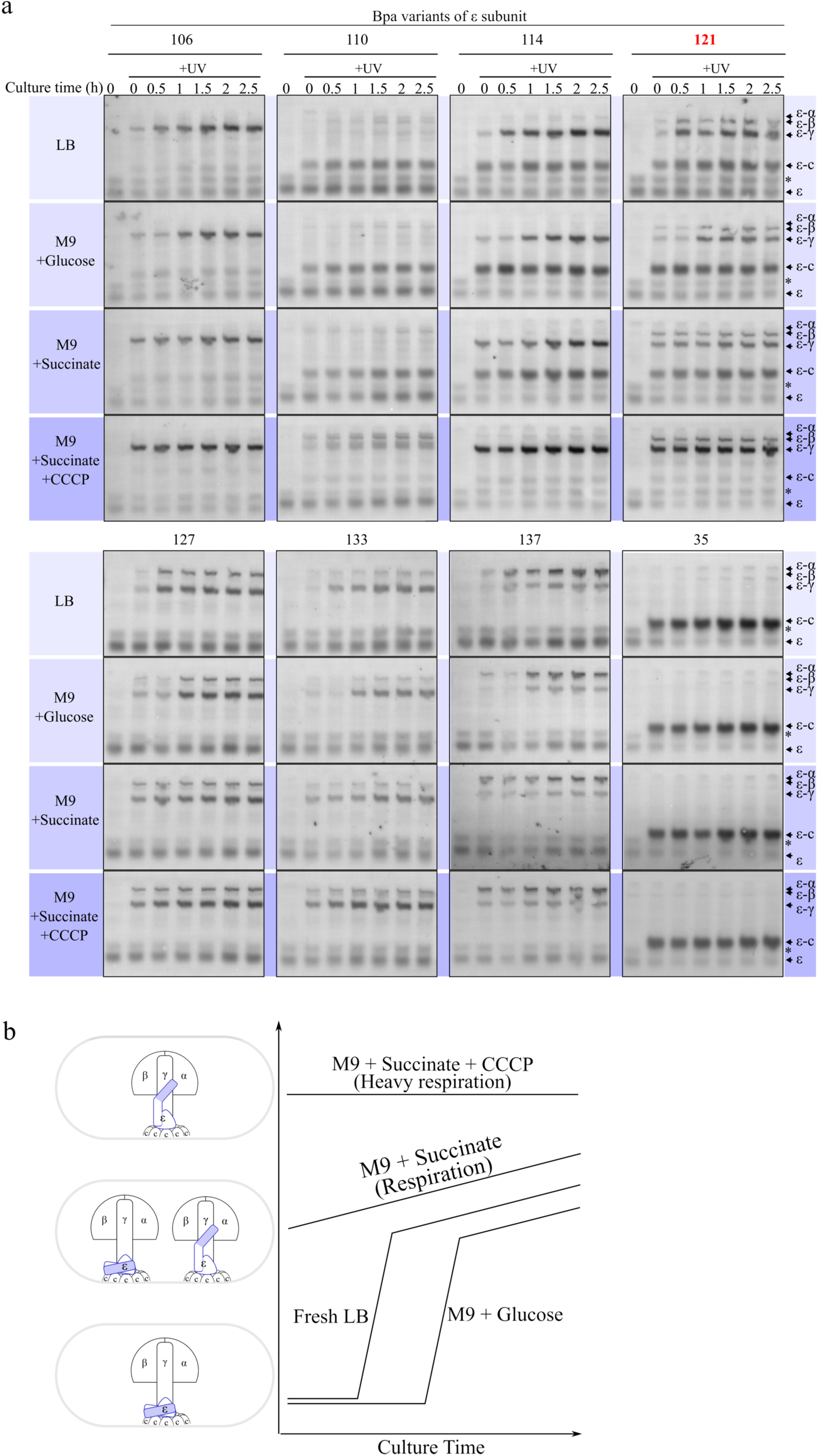
The ε-subunit of the ATP synthase spontaneously switches from a “non-inserted” state to the “inserted” state after the cells are growing for certain durations either in fresh LB rich medium or in glucose-containing M9 minimal medium. (**a**). Blotting results for detecting the photocrosslinked products of seven selected Bpa variants of the C-terminal helix of the ε-subunit after each type of the cell was cultured in the indicated medium for the indicated durations. The variant having a Bpa introduced at residue position 35 (in the N-terminal domain) was analyzed here for comparison. The protein samples were probed with alkaline phosphatase-conjugated streptavidin against the ε-subunit (as in Fig. 4a). Positions of the ε monomer or crosslinked ε products are indicated on the right of the gel; again, * indicates a non-specific band. (**b**). A schematic summary of the time-dependent existence of “inserted” and “non-inserted” states for the ε-subunit in cells cultured in the indicated media, as indicated by the crosslinking results displayed in (**a**).

Notably, in correlation to the increase of the level of the “inserted” state, the level of the “non-inserted” state as reflected by the formation of crosslinked products between the C-terminal helix of the ε-subunit and the c-subunit was not found to be decreased, being either unaltered or even increased along the culturing process of the cells in all these three media (**Fig. 5a**). These observations implies the likely existence of another “non-inserted” form besides that represented by the interaction between the C-terminal helix of the ε-subunit and the c-subunit. It is conceivable that this new “non-inserted” form might be represented by the previously described half-extended state of the C-terminal domain of the ε-subunit (Sobti et al., 2019). Remarkably and interestingly, when the cells were cultured in the M9 minimal medium containing CCCP (at a concentration of 10 μM) in addition to the succinate, the “inserted” state of the C-terminal helix of the ε-subunit became the most predominant at the 0 time point of culturing. More importantly, the “non-inserted” state as represented by an interaction of the C-terminal helix of the c-subunit became only slightly detectable (the “M9 + succinate + CCCP” panels in **Fig. 5a**; best indicated by the 121-Bpa and 114-Bpa variants). at this point, it is worthwhile to note that we also demonstrated that the cells were able to grow effectively when placed in this CCCP- and succinate-containing M9 minimal medium, though at a rate slower than when the cells were cultured in the M9 medium containing only succinate in the absence of CCCP (the data are shown in **Fig. 7a**, the blue and black pairs of curves).

Collectively, these observations suggest that it is the “inserted” state, rather than the “non-inserted” state, seems to be important in keeping the ATP synthase in a functional state under such unfavorable environmental conditions as growing in the spent medium or in CCCP-containing M9 minimal medium. These observations further supported our conclusion that the “inserted” and “inserted” states exist as a dynamic equilibrium, spontaneously shifting to one side or the other depending on the environmental conditions.

### Hypothesis: the “inserted” state of the ε-subunit that is predominant under unfavorable physiological conditions strengthens the ATP synthesis function of the ATP synthase

These above described observations, achieved by examining the interaction patterns of the C-terminal helix of the ε-subunit in living cells, with the data mainly displayed in **Figs 4** and **5**, clearly demonstrate that the ε-subunit is able to switch to either the “inserted” or the “non-inserted” state in suiting to the particular environmental conditions under which the cells are living. Specifically, the “inserted” state is predominant in cells living under such unfavorable conditions as in the overnight-cultured spent LB rich medium or in CCCP-containing M9 minimal medium, but is minor in cells living under such favorable conditions as in fresh LB rich medium or glucose-containing M9 minimal medium (**Fig. 4**). Moreover, the “non-inserted” state would be converted to the “inserted” state after the cells are cultured in the fresh LB rich medium or in glucose-containing M9 minimal medium for a period of time (**Fig. 5**), apparently reflecting a change of their growth condition from the initially favorable one to the subsequently less favorable one.

In light of our observation that the “inserted” state of the ε-subunit seems to allow the bacterial cells to grow and divide, though in a somehow less efficient way, under the unfavorable conditions (to be shown below in **Fig. 7a**). We hypothesize that the “inserted” state of the ε-subunit strengthens the ATP synthesis function of the ATP synthase, hence enabling the cells to grow and divide under an unfavorable (or less favorable) physiological condition, whereas the “non-inserted” state represents a non-strengthened form of the ATP synthase that exists mainly in cells living under a favorable physiological condition.

### Certain mutations of the critical amino acid residue I125 in the ATP synthase ε-subunit lead to a severely defective formation of the “inserted” state

We next attempted to provide functional evidences that support the hypothesis we proposed above. In this regard, it needs to be pointed out that a growth phenotype analysis of mutant cells in which the C-terminal helix of the ε-subunit was completely truncated engendered the authors to conclude that this C-terminal helix is dispensable for growth and survival of the *Escherichia coli* bacterial cells (Taniguchi et al., 2011). Nevertheless, for the purpose of elucidating the biological role of the “inserted” and “non-inserted” states of the ε-subunit, such truncation studies might be considered hardly proper, since none of the two states was maintained. We reasoned that it would be difficult for us to put our hypothesis to a test unless we could generate and identify a type of mutants for which the formation of the “inserted” state is severely defective, whereas the formation of the “non-inserted” state remains largely unaffected (or strengthened).

As an attempt of identifying such mutant forms of the ε-subunit, we decided to perform a screening on all the 19 possible replacement mutants of residue I125, which was chosen because of its apparent strong interaction with the β-subunit, as reflected by the crosslinking results displayed in **Figs. 1b** and **4a**. This in turn suggested that I125 might play an important structural role in forming the “inserted” state of the ε-subunit in living cells. However, this screening would not be feasible unless we could effectively monitor the conformational state of all these 19 mutant ε-subunits. In this regard, photocrosslinking mediated by the Bpa introduced at the position of residue 121, which had been demonstrated to be able to either interact with the α/β/γ-subunits thus indicating the existence of the “inserted” state or with the c-subunit indicating the existence of the “non-inserted” state in living cells (**Figs. 1b** and **4a**), seems to serve as an ideal and unique way for this monitoring purpose. We thus generated these 19 mutant ε-subunits by performing genome-targeted site-directed mutagenesis on the codon of residue I125 using cells producing the 121-Bpa variant of the ε-subunit (i.e., each ε-subunit variant contained two replaced residues). Additionally, we examined the conformational state of all the 19 mutants generated on I125 by treating the mutant cells for ~10 min with exactly the same set of experimental conditions that we described above (**Fig. 5a**).

The blotting results for detecting the photocrosslinked products, as displayed in **Fig. 6**, revealed that, to our delight, among these 19 mutants of the ε-subunit, eight of them did produce a significantly decreased level of crosslinked products with the α/β/γ-subunits, reflecting a severe defect in forming the “inserted” state, almost under all the tested experimental conditions (the middle column). Moreover, the interactions between the C-terminal helix of these eight ε variants with the c-subunit seemed to be hardly affected, i.e., occurring at a level largely comparable with that of the original unmutated I125-ε-subunit (top left). Out of these eight mutants, the I125K variant (red colored in **Fig. 6**) exhibited the most severe defect. The rest eleven I125 variants could be roughly categorized into the following three groups. Four of them (left column) produced a level of either the “inserted” or the “non-inserted” state highly comparable with the unmutated I125-ε-subunit (which was also included in the left column); five of them (right column, top part), though produced a largely unaltered level of the “inserted” state, became incapable of interacting with the c-subunit; two of them (right column, bottom part) became incapable of interacting with either the α/β/γ-subunits (thus unable to form the “inserted” state) or the c-subunit (thus existed possibly as the other “non-inserted” state that we described above).

**Figure 6.**
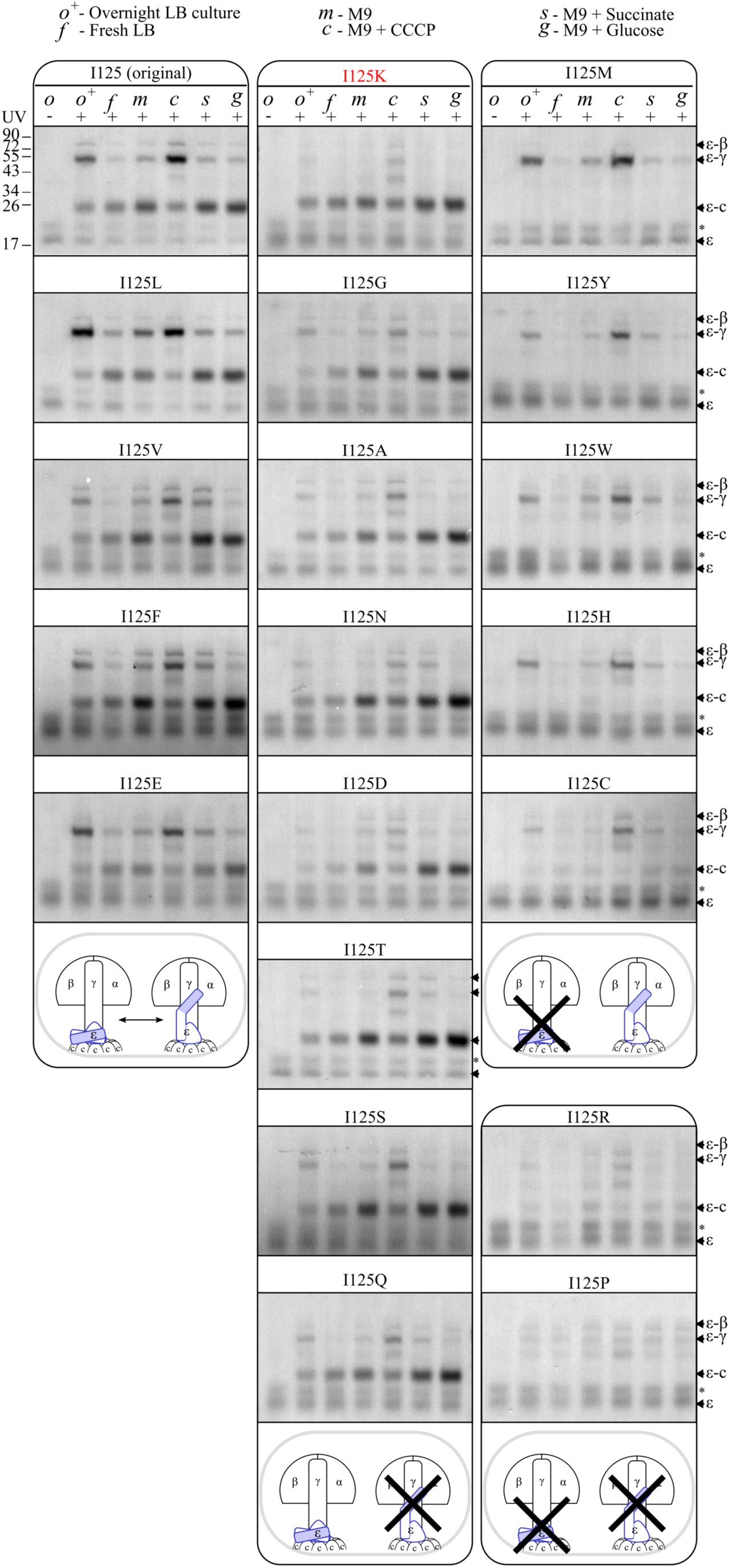
Certain replacements of the critical amino acid residue I125 of the ε-subunit engender the formation of the “inserted” state to be defective. Blotting results for detecting the photocrosslinked products formed (via Bpa introduced at residue position 121) by the 19 replacement variants generated at residue I125 after the modified cells were treated by the same solutions described in Fig. 4a. Again, all the samples were probed with alkaline phosphatase-conjugated streptavidin against the ε-subunit. Positions of the ε monomer and crosslinked ε products are indicated on the right of the gel; * indicates a non-specific band. The major states detected for the mutants of the ε-subunit are summarized at the bottom of each group (with a cross placed on the state that became hardly detectable).

In sum, these data suggest that residue I125 plays a key role for the C-terminal helix of the ε-subunit to interact with either the α/β/γ-subunits in forming the “inserted” state or to interact with the c-subunit in forming one “non-inserted” state. More importantly, we identified the ε-I125K variant as one who became severely defective in forming the “inserted” state, but was hardly affected in forming the “non-inserted” state. This ε-I125K mutant cell was exactly what we wanted to generate and was thus chosen for further phenotype analyses.

### The “inserted” state of the ε-subunit provides a moderate but significant advantage for the bacterial cells to grow under unfavorable physiological conditions

As a way to provide supporting evidence to our hypothesis, we next examined the growth phenotype of this ε-I125K variant (in which Bpa was no longer introduced at residue position 121). In particular, we first compared the growth rate of the ε-I125K mutant and the wild type bacterial cells that were cultured in the M9 minimal medium containing succinate and CCCP (at 10 uM), in which the mutant cells would only form a very low level of the “inserted” state (as demonstrated above, **Fig. 6**) whereas the wild type ε would exist mainly as the “inserted” state (also as demonstrated above, **Fig. 5**). The growth curves presented in **Fig. 7a** demonstrated the following. First, both the ε-I125K mutant and the wild type cells were able to grow in this medium (the blue pair of curves), though not as efficiently as they did in the M9 minimal medium containing only succinate but not CCCP (the black pair of curves). Second, there existed only a moderate decrease of the growth rate and only at the later stages of culturing for the ε-I125K mutant cells in comparison with the wild type cells (the blue pair of curves).

**Figure 7.**
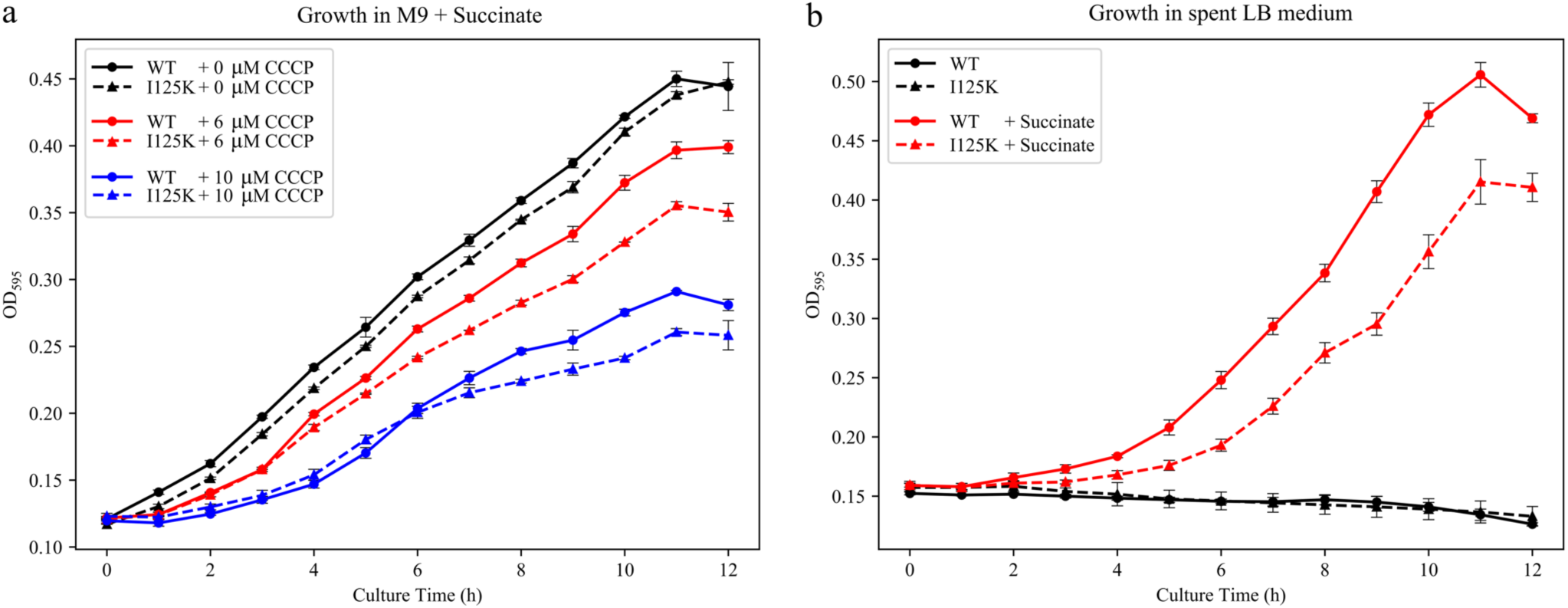
The ε-I125K mutant cell that is severely defective in forming the “inserted” state exhibits a moderate but significant disadvantage in growing under unfavorable conditions. Shown are the paired growth curves of the bacterial cells that respectively possessed the mutant ε-I125K (dash lines) and wild type ε-(solid lines) subunits and were cultured in the indicated succinate-containing M9 minimal medium with an addition of the indicated concentrations of CCCP (**a**), or in the succinate-supplemented spent LB rich medium (**b**). The growth curves were generated by plotting the optical density at 595 nm (OD_595_) against the culturing time points. The standard deviation values were calculated based on the measurements of three repeated experiments.

We reasoned that adding CCCP at 10 uM might have produced a condition that was too unfavorable for the wild type ATP synthase to cope with. We thus repeated these experiments to make the conditions more favorable (or less unfavorable) by decreasing the CCCP concentration to 6 uM. Interestingly, consistent to our expectation, we did observe a larger degree of decrease in the growth rate for the ε-I125K mutant cells in comparison with the wild type cells (the red pair of curves). This implied that the formation of the “inserted” state provides an advantage on cellular growth under a moderately unfavorable rather than a severely unfavorable condition. Apparently, such a rather moderate difference in the growth rate would be difficult to distinguish simply by examining the mutant cells in which the C-terminal helix of the ε-subunit was completely removed (Taniguchi et al., 2011).

This observed growth defect of the ε-I125K mutant cells prompted us to find out whether the same could be demonstrated when the cells were cultured under a fully natural condition, i.e., in the absence of the exogenously added uncoupler CCCP. To this end, we recalled the fact that the crosslinking pattern of the C-terminal helix of the ε-subunit in cells overnight cultured in LB rich medium (the *o^+^* lanes in **Fig. 4a**) was highly comparable to that detected when the cells were treated in the CCCP-containing M9 minimal medium (the *c* lanes in **Fig. 4a**). This somehow suggested that the wild type ε might predominantly have existed as the “inserted” state in the overnight cultured LB rich medium due to the action of certain CCCP-like chemical agent, which was produced by the physiological activities of the cells themselves. In this regard, it is worth pointing out that the indole molecule was reported to accumulate in the medium after the bacterial cells were cultured in LB rich medium and has been described as a naturally produced CCCP-like respiration chain uncoupler (Chimerel et al., 2012; Gaimster et al., 2014). Furthermore, our observation that the wild type ε-subunit mainly existed as the “inserted” state in the M9 medium containing both CCCP and succinate (**Fig. 4**, the *c* lanes) would implicate the same to occur in the spent LB rich medium in which succinate is added.

We hence compared the growth rate of the ε-I125K mutant and the wild type cells in the spent LB rich medium containing succinate (in which the former would predominantly exists as the “non-inserted state while the latter as the “inserted” state). Interestingly, the degree of growth rate for the ε-I125K mutant cells in comparison with the wild type cells was found to be even larger in this medium, indicating a more drastic disadvantage for the mutant cells to grow under this somewhat natural growth condition (the red pair of curves in **Fig. 7b**). As expected, neither the wild type nor the mutant bacterial cells were able to grow in the spent LB culture medium when succinate was not added as a carbon source (the black pair of curves in **Fig. 7b**). Again, this observation was in full agreement with our hypothesis that the “inserted” state of the ε-subunit has been evolved for the cells to supply enough energy to cope with a moderately unfavorable, rather than a severely unfavorable condition. Evidently, such a moderate growth advantage would accumulate to a major one over a long duration and allow the bacterial cells to effectively grow under an unfavorable condition.

Further, we also spent much effort in measuring the intracellular concentration of the ATP molecule present in the ε-I125K mutant and wild type cells that were cultured in all the above-described culture media (**Figs. 7a** and **7b**). Nevertheless, we failed to reveal any ATP concentration differences of statistical significance. In other words, the measured differences in ATP concentration between the ε-I125K mutant and wild type cells were found to be in the same range as the variations we obtained between the repeating experiments. These results apparently implied that the differences in ATP levels between the mutant and wild type cells are most likely very minor even under these unfavorable experimental conditions despite of the moderate differences in growth rate.

## Discussion

This study was conducted for the purpose of deciphering the physiological relevance of *in vitro* biochemical observations on the ATP synthase ε-subunit, being a common issue in current protein researches. In particular, we attempted to define the physiological context under which the “inserted” state of the ε-subunit of the bacterial ATP synthase might be formed to a significant level and also to elucidate the biological role such a state might play. We addressed these issues by performing systematic high-throughput analyses of the photocrosslinked products formed via the unnatural amino acid Bpa genetically incorporated at a large number or residue positions, which allowed us to directly reveal the dynamic interaction patterns of the ε-subunit in living cells that were treated (for a very short period of time) or cultured (for several hours) under a variety of different experimental conditions. These analyses were made possible as a result of our development of two new techniques, the scarless genome-targeted site-directed mutagenesis technique for introducing the unnatural amino acid Bpa by mutating the endogenous gene encoding the ε-subunit (**Figs. 1a** and **S1a**), and the high-throughput polyacrylamide gel electrophoresis (HT-PAGE) technique for performing electrophoresis and blotting analyses of up to 384 protein samples on one single gel (**Fig. 2**).

By employing these new techniques, we revealed for the first time that the ε-subunit is able to predominantly exist as the “inserted” state under such common conditions as in the spent LB rich medium. We further demonstrated that the ε-subunit is able to instantly and reversibly switch between the “inserted” and the “non-inserted” states in responding to the fluctuating environmental conditions (**Figs. 3–5**). Importantly, by identifying the ε-I125K mutant as one that is severely defective in forming the “inserted” state, i.e., largely existing as the “non-inserted” state (**Fig. 6**), we were able to demonstrate that the formation of the “inserted” state provides a moderate but significant growth advantage for the cells to live under unfavorable conditions (**Fig. 7**). Such a highly reversible fine-tuning mechanism that will empower the ATP synthase and thus allow the cells to grow and divide under unfavorable conditions. Such a mechanism for the function modulation of the ATP synthase apparently occurs in a way analogous to gear shifting in the power transmission system of an automobile, as schematically illustrated in **Fig. 8**. The major features of this mechanism might be described as such that the ε-subunit acts as an efficient molecular switch that shifts to the low-gear “inserted” sate to strengthen the ATP synthesis function of the ATP synthase to supply enough energy to the cells in coping with unfavorable physiological conditions, whereas to immediately shift back to the high-gear “non-inserted” non-strengthened state when the environmental conditions resumes to favorable and functional strengthening of the ATP synthase becomes unneeded.

**Figure 8.**
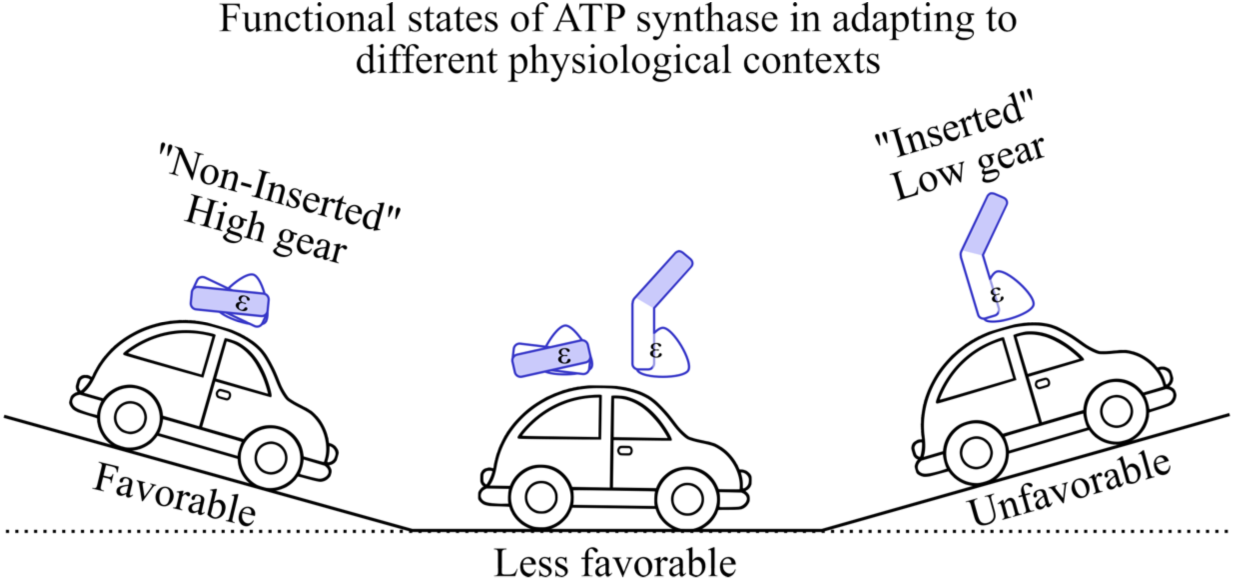
Schematic diagram illustrating the fine-tuning regulation mechanism of the ATP synthase functioning as contributed by the formation of the “inserted” low-gear and “non-inserted” high-gear states of the ε-subunit. As in the power transmission system of an automobile, the “inserted” low-gear state of the ATP synthase represents a functionally strengthened form that exists in cells living under an unfavorable environmental condition, whereas the “non-inserted” high-gear state represents a non-strengthened form of the ATP synthase that exists in cells living under a favorable environmental condition. An ATP synthase complex is able to switch between these two states in an instant and reversible manner allowing the cells to effectively respond and adapt to the frequent fluctuations of their environmental conditions.

It is conceivable that the ATP synthesis capacity of the ATP synthase in the low-gear “inserted” state is not as strong as when the ε-subunit is in the high-gear “non-inserted” state. But this represents an effort of the cell to keep its ATP synthase in the best functional state in coping with the unfavorable, cellular environment, which may or may not last long. This “inserted” state, although only strengthens the function of the ATP synthase to a moderate degree, it is formed quickly and reversibly and thus allow the cells to efficiently adapt to their fluctuating environmental conditions. This would allow the ATP synthase to keep functioning despite of the presence of a certain level of natural uncouplers, which are believed to be produced under common metabolic processes of the cells (Chimerel et al., 2012; Gaimster et al., 2014), or other have yet to be identified unfavorable conditions. It would be interesting to find out whether the ε-subunit in the chloroplast ATP synthase also functions in a similar manner. Notably, *in vitro* structure analysis indicated that similar “inserted” and “non-inserted” states seem to be formed by the inhibitor protein IF_1_ present in mitochondria (Walker, 2013). Whether IF_1_ also function via a similar gear-shifting mechanism in regulating the function of the ATP synthase in mitochondria is worth further investigation. Evidently, such a gear-shifting type of fine-tuning mechanism would be an efficient way for many enzymes to modulate their function, thus to maintain the homeostatic state of the cells/organisms in responding to the frequently occurring fluctuations, rather than dramatic changes, in their environmental conditions. The evolution of such a quick and reversible mechanism of functional modulation might represent a vital aspect for the lives of cells, at least for such single-cell organisms as bacteria.

Our living-cell observations reported here are largely in agreement with multiple *in vitro* findings on the structure features of the C-terminal domain of the ε-subunit. For example, first, our findings are in agreement with the hypothesis proposing that the “up” conformation (corresponding to the “inserted” state) might inhibit the ATP hydrolysis activity of the ATP synthase (Tsunoda et al., 2001). This could be viewed as one way to strengthen the function of the enzyme complex under unfavorable conditions. Second, our observations are not in disagreement with the structural revelation that the C-terminal domain of the ε-subunit formed the “up” conformation in the absence of ATP but the “down” conformation (corresponding to our “non-inserted” state here) in the presence of ATP (Yagi et al., 2007; Sobti et al., 2019), although it is neither directly supported by our observations. Third, our data provided living-cell evidences to support the existence of a half-extended conformation for the C-terminal domain of the ε-subunit, which forms neither the “up” nor the “down” (as represented by an interaction between the C-terminal helix of the ε-subunit and the c-subunit) conformation (Sobti et al., 2019). This half-extended conformation correlates to the one in which the C-terminal helix interacts with neither the α/β-subunits nor the c-subunit in the ATP synthase (but likely to interact with the γ-subunit). Fourth, our observations support the view that the formation of the “inserted” or “up” state is related to the energy state of the cell (Feniouk et al., 2006; Walker, 2013), but we are not certain whether the ATP synthase responds to the level of ATP, or of proton motif force, or of both, or even of other factors.

The two techniques that we developed here, the scarless genome-targeted site-directed mutagenesis and the HT-PAGE, could have potential applications in many other areas of protein studies for and beyond unnatural amino acid-mediated photocrosslinking analysis. The scarless genome-targeted site-directed mutagenesis technique would allow one to replace codons of any protein-encoding genes in the authentic genome context, such that the timing and quantity of production for the modified proteins will be identical to that of the wild type protein, which has often been highly desired in learning about the function of a particular protein. by contrast, the target proteins have been conventionally modified by manipulating their encoding genes carried by an exogenously introduced DNA expression vectors (based on plasmids or virus), such that the timing and quantity of the production of the modified protein would be significantly different from that of the endogenous wild type protein, a situation highly undesirable but difficult to avoid in many studies.

The HT-PAGE technique provides a powerful way to analyze a large number of protein samples via exactly the same operation procedure of electrophoresis and blotting analysis, which would thus allow a semi-quantitative comparison of all the protein bands, either after being directly stained with a dye (like Coomassie Brilliant Blue) or after being blotted with a specific probe (such as an antibody), as often highly desired in many proteomics studies that often explore the quantitative changes of particular proteins under different conditions (like from normal to cancer cells). By contrast, the currently employed vertical electrophoresis system (either one- or two-dimensional) only allow the examination of a limited number of protein samples on each gel, i.e., for a large number of protein samples to be analyzed, multiple different gels have to be operated in parallel, which would unavoidably generate operational variations in the results from one gel to another and thus problematic if one wants to perform a semi-quantitative comparison of one particular protein band across all the gels. In our case, this system also provided us an extraordinary way to self-validate the crosslinking pattern identified based on one Bpa variant by confirming it with all the other Bpa variants introduced in the same domain of the protein. This will undoubtedly prove to be a powerful approach in analyzing dynamic protein-protein interactions occurring in living cells, for which such self-validations would be essential for making any reliable conclusions.

In sum, our high-throughput polyacrylamide gel electrophoresis (HT-PAGE) and blotting analyses in combination with scarless genome-targeted site-directed mutagenesis allowed us to systematically examine the protein-protein interactions of the ATP synthase ε-subunit in living cells placed under a great variety of experimental conditions. These studies enabled us to unveil the long-sought physiological relevance of the alternative “inserted” and “non-inserted” quaternary structures of the ATP synthase ε-subunit that have been characterized mainly based on *in vitro* studies. Our approach provided a new strategy for uncovering the physiologically relevance for the *in vitro* biochemical observations on proteins, an outstanding and critical issue in the current field of molecular biology.

## Materials and Methods

### Bacterial strains

Listed in **Table S1** are the genotypes of all the *E. coli* bacterial strains that were used in this work, all being derived from the *E. coli* strain BW25113, whose genotype is as follows: F-, DE(araD-araB)567, lacZ4787(del)::rrnB-3, LAM−, rph-1, DE(rhaD-rhaB) 568, hsdR514 (Grenier et al., 2014).

### Genome-targeted site-directed mutagenesis

All genome modifications were performed via two rounds of homologous recombination using the λ-red genomic recombination system (Lee et al., 2009). As shown in **Fig. S1a**, the first round of recombination would generate the Δε strain from the *E. coli* strain LY928, after the gene encoding the ε-subunit was replaced by the kanamycin resistance gene (*kana*^r^). Then in a second round of recombination, the Avi-ε strain (designated as the wild type strain for convenience) and all the strains producing the Bpa variants of the Avi-ε-subunit were generated via a scarless replacement of the *kana*^r^ in the Δε strain by a gene encoding Avi-ε or a variant of Avi-ε (i.e., having a TAG replacement at a selected codon position).

A successful codon replacement was indicated and selected via the production of a functional ATP synthase, which is needed in supporting the normal growth (i.e., with the formation of large normal colonies) of the recombined variant bacterial strain in solid agar-containing LB rich medium. In contrary, a failure of the second round of the recombination would be indicated by having the cells to grow in a defective way, i.e., only to form small colonies as the Δε strain (a representative result is displayed at the bottom of **Fig. S1a**). In addition, all the genome modifications were eventually confirmed by DNA sequencing.

The functional and structural status of each Bpa-containing ε variant were examined by directly examinig whether the resulted bacterial strain was able to grow in succinate-containing M9 minimal medium (47.7 mM Na_2_HPO_4_, 22.0 mM KH_2_PO_4_, 8.55 mM NaCl, 18.7 mM NH_4_Cl, 1 mM MgSO_4_, 0.2% succinate), such that the ε variants that did not allow the formation of a functional ATP synthase would not engender the bacterial cells to grow (as shown in **Fig. S1b**).

### Conventional cell culturing, protein photocrosslinking and Western blotting

The cells were cultured overnight at 37°C in Bpa-containing (at 200 uM; F2800, Bachem) sterilized LB liquid rich medium (10 g/l tryptone, 5 g/l yeast extract, and 5 g/l NaCl), with an agitation of ~ 260 r.p.m. The cells were then irradiated with UV light (365 nm) for ~10 min at room temperature using a Hoefer UVC 500 Crosslinker (Amersham Biosciences, USA), collected by centrifugation at 13,000 × g, resuspended in the sample buffer, boiled and resolved by tris-tricine SDS-PAGE. The Avi-ε-subunit and its photocrosslinked products were probed by alkaline phosphatase (AP)-conjugated streptavidin (which will bind to the biotin attached to the Avi-tag). The α-, β- and γ-subunits, as well as the their photocrosslinked products formed with the Bpa variants of the ε-subunit were respectively probed against their particular antisera (laboratory stock). The protein bands visualized on the PVDF membranes were scanned and processed using the GNU image manipulation program (**Figs. 1b** and **2**).

### Developing a high-throughput SDS polyacrylamide gel electrophoresis (HT-PAGE) technique

Before developing our HT-PAGE, we noticed that Barbieri *et al* reported the development of a horizontal sodium dodecyl sulfate polyacrylamide gel electrophoresis (SDS-PAGE) technique that could be applied to analyze both protein and DNA samples. However, the authors did not consider to develop it into a high-throughput technique (Izzo et al., 2006). We thus prepared a horizontal polyacrylamide gel between an acrylic plate that contains one row of 15 protruding teeth of 3 × 1 × 1 mm (particularly made for us by the supplier, from Taobao.com) and a microscopic glass slide (75 × 25 mm) using tris-borate (89 mM) as the gel buffer, as adapted from their report (Izzo et al., 2006). We tested this system by performing an electrophoresis and blotting analysis on the photocrosslinked products of 12 of the Bpa variants of the ε-subunit. However, to our disappointment, we found that the resolution was too low and did not allow us to distinguish the protein bands representing the ε-subunit, the nonspecific protein and the ε-c photocrosslinked product (refer to **Fig. 1b**). In an attempt to improve the resolution, we randomly tested a variety of electrophoresis buffers commonly that have been used for DNA and protein electrophoresis, including, for example, the Tris-Borate, Tris-Acetate, Tris-Glycine, Bistris-MOPS and Tris-Tricine (all at 100 mM). The blotting results demonstrated that, for reasons unclear to us, the buffer composition did exhibit a profound effect on the resolution of the gels, while the Tris-Acetate buffer system gave the poorest resolution, the Tris-Tricine buffer gave a greatly improved resolution.

As the polyacrylamide composition (bisacrylamide/total acrylamide or *c* = 5%) described by Barbieri et al (Izzo et al., 2006) is different from the polyacrylamide composition (c=2.6%) commonly used for protein analysis, we next evaluated the effect of different polyacrylamide compositions (with a *c* value of 2.5%, 3%, 4%, 5%, 6% or 7%) on gel resolution. Again, the blotting results demonstrated that the resolution varied significantly with different acrylamide compositions, either a low or a high *c* value would sabotage the resolution, only with the c = 6% ratio gave us the highest resolution.

Taken together, after these screening operations on both gel compositions and buffers, we achieved an initial success in developing a submerged horizontal SDS-PAGE system via an utilization of the 100 mM tris-tricine solution as the gel preparation and electrophoresis buffer, as well as the preparation of the gel as one containing 11% acrylamide and c=6%. This combination of conditions allowed us to successfully separate the ε-subunit and all its photocrosslinked products (with a molecular mass range of 15 -75 kDa). We designate this as our stage 1 submerged horizontal SDS-PAGE system, with the size of each lane to be 3 × 20 mm (Stage 1 in **Fig. 2**).

Towards the goal of developing HT-PAGE, we next had to find a way to increase the number of protein samples that could be analyzed per gel. In this regard, we modified the acrylic plate such that it now contained one additional row of protruding teeth, which accordingly led to a decrease of the migration length by half for each protein sample, from 20 mm to 10 mm. The blotting results demonstrated a minor but acceptable decrease in the resolution, although the number of protein samples analyzable increased from 15 to 30. We designate this as our stage 2 submerged horizontal SDS-PAGE system, with the size of each lane to be decreased to 3 × 10 mm (Stage 2 in **Fig. 2**).

To further double the number of protein samples that could be analyzed per gel (from 30 to 60), we next decided to decrease the size of each protruding teeth as well as the space between the neighboring teeth, resulting in the width per lane to be reduced to 1 mm (rather than 3 mm as in Stages 1 and 2). This modification of the equipment led to a decrease of the volume of the samples loaded per well from 3 ul (in Stages 1 and 2) to 1 ul, as well as a corresponding increase of the time needed to load all the 60 samples. Our initial results revealed a new problem, i.e., the protein samples loaded at an earlier time point would get lost to a significant degree. To overcome this, we reasoned that an increase in the viscosity of the electrophoresis system might reduce the diffusion of the protein samples. To alleviate this problem, we decided to add glycerol in the running and loading buffer, as well as in the polyacrylamide gel. Again, after testing a variety of different concentrations, we found that glycerol at a concentration of 10% (w/v) prevented the diffusion problem of the protein samples, as well as with a resolution largely comparable to what we achieved in Stages 1 and 2. We designate this as our stage 3 submerged horizontal SDS-PAGE system, with the size of each lane to be decreased from 3 × 10 mm to 1 × 10 mm (Stage 3 in **Fig. 2**).

Data obtained in stage 3 revealed that we might be able to further increase the number of protein samples that could be analyzed per gel by increasing the overall size of the gel, provided that we could load all the samples within about 30 min. In this regard, it was apparent to us that we could speed up the sample loading process by using the commercially available multi-channel pipette. Towards this goal, we decided to further slightly reduce the lane length for each sample to fit the 8 mm space between each channel of the eight-channel pipette. In line with this new design, we accordingly prepared the acrylic plate as consisting 8 rows of teeth, with each row made of 48 teeth (thus allowing a total of 384 samples to be analyzed per single gel). Unfortunately, in testing this new gel system with the buffer that worked effectively at stage 3, we failed to achieve a satisfactory resolution (judging by its capacity to resolve the three closely spaced protein bands we mentioned above). Inspired by what we achieved in stage 1, we performed a similar random screening for proper buffers, wishing for similar luck in finding an optimal one. For this purpose, we surveyed a variety of buffers that were prepared as a combination of various anions (Tris, triethanolamine, Bis-tris, and imidazole) and cations (Glycine, Tricine, MOPS, and HEPES). To our satisfaction, we found that the triethanolamine-HEPES combination buffer allowed us to achieve a resolution high enough to clearly distinguish even the closely spaced protein bands. We designate this as our stage 4 (i.e., the final) submerged horizontal SDS-PAGE system, with the size of each lane to be slightly decreased from 1 × 10 mm to 1 × 8 mm (Stage 4 in **Fig. 2**).

After going through these four stages of improvements, we successfully developed a 384-lane HT-PAGE system. In sum, the operation of this system requires the following components: 1) the 30% acrylamide gel stock solution (T=30%, C=6%); 2) the 5 X gel and running buffer solution (250 mM Triethanolamine, 250 mM HEPES, 50% glycerol (v/v), 0.5% SDS (w/v); 3) the 2 X loading buffer (200 mM tris-phosphate buffer pH 6.8, 4% SDS (w/v), 20% glycerol (v/v), 200 mM DTT, 10% Glucomannan (w/v), bromophenol blue 0.008% (w/v). Specifically, the 384-lane gel could be prepared according to the following protocol. First, the acrylamide gel stock solution was mixed with the 5 X gel and running buffer and water to make an 11% acrylamide solution. Second, after adding the polymerization initiator, the gel solution was injected to the space between the protruding teeth-bearing acrylic plate and the covering glass plate. Third, the polymerization initiator could be ammonium persulfide, or more preferably omnirad 2959 (IGM, former BASF) which would initiates the polymerization process only after an UV light (365 nm) irradiation. Fourth, after polymerization, the gel sticking onto the glass plate could now be placed in a common horizontal electrophoresis device for performing the electrophoresis. Such a prepared gel could be stored at 4°C for 1 year without a significant loss of resolving power.

### High-throughput cell treatments and *in vivo* photocrosslinking

The high-throughput pipeline for culturing, treatments and photocrosslinking of the bacterial cells, as well as protein electrophoresis and Western blotting are schematically illustrated in **Fig. 3**. First, each bacterial strain (including all the 36 Bpa variants of the C-terminal helix of the ε-subunit, and all the 20 variants of residue I125 generated with the 121-Bpa ε-subunit) was individually cultured in a well harboring 1 ml Bpa-containing LB rich medium via the utilization of a 96-deep-well plate. After the wells on the plate was covered with a sterile breathable sealing film (BF-400-S, Axygen), the cells were then incubated overnight at 37°C with an agitation at 500 r.p.m. Second, the cultured cells were subsequently aspirated by using an eight-channel pipette and dispersed into 7 equal aliquots (40 ul each) using a 96-well PCR plate, centrifuged at 2700g for 15 min to remove the medium (as the supernatant). The remaining cells (as the pellet) were afterwards resuspended in the following 7 solutions: the spent LB medium (two sets, one with and the other without UV irradiation), the fresh LB medium, the M9 minimal medium (47.7 mM Na_2_HPO_4_, 22.0 mM KH_2_PO_4_, 8.55 mM NaCl, 18.7 mM NH_4_Cl), the M9 minimal medium containing 10 uM carbonyl cyanide 3-chlorophenylhydrazone (aka. CCCP; C2759, Sigma-Aldrich), the M9 minimal medium containing 0.2 % (w/v) succinate, and the M9 minimal medium containing 0.2 % (w/v) glucose. Third, immediately after the treatments (~ 10 min), the 96-well PCR plates were subjected to UV irradiation (365 nm) for ~10 min, followed by an addition of 40 ul of the 2 X loading buffer and a heating at 95°C for 5 min to complete the sample preparation process. Fourth, the samples were then transferred to a 384-well plate according to a predesigned loading plan, loaded onto the 384-well polyacrylamide gel by using the 8-channel pipette. After electrophoresis, the gel was blotted and probed with AP-conjugated streptavidin against the Avi-ε-subunit of the ATP synthase (**Figs. 4** and **6**).

### High throughput *in vivo* photocrosslinking analysis of the selected ε variants after the cells were cultured in various media for 30-min intervals

The cells were all first cultured overnight in a 96-deep-well plate (with each well containing 1 ml Bpa-containing LB rich medium). The cultured cells were then each equally aliquoted to 4 portions (150 ul each) and transferred to a new 96-well PCR plate, centrifuged at 2700 g for 15 min to remove medium. The resulted pellet cells were the resuspended in four culture media: the fresh LB rich medium, as well as the M9 minimal medium (1 mM MgSO_4_, 1 mM thiamine, 0.2 % (w/v) casamino acids) containing 0.2 % glucose, or 0.2 % succinate, or 0.2 % succinate and 10 uM CCCP. The resuspended cells were then transferred to a 96-well plate, covered with sterile breathable sealing film and cultured at 37 °C with agitation. At the indicated time points (0, 30, 60, 90, 120, and 150 min), an aliquot (of 10 ul) of each inoculated culture was removed, irradiated with UV light (365 nm). After all samples were collected, they were subjected to HT-PAGE/western blotting analysis (**Fig. 5**).

### Growth analysis of the wild type and ε-I125 variant strains

The wild type (WT) and ε-I125 variant strains were cultured overnight at 37 °C in LB rich medium in test tubes with an agitation of ~260 r.p.m. The cultured cells were then each dispersed into 5 aliquots (40 ul each) before respectively added into a 600 ul of the following five culture media: the M9 minimal medium containing succinate (containing 0.2 % (w/v) succinate, 1 mM MgSO_4_, 1 mM thiamine, 0.2 % (w/v) casamino acids in addition to the M9 buffer); the M9 minimal medium containing succinate and 10 uM CCCP; the M9 medium containing succinate and 6 uM CCCP; the spent LB rich medium; the spent LB rich medium containing 0.2 % succinate. Each of these cell samples was then divided into three equal parts and transferred to a 96-well plate, covered with sterile breathable sealing film, cultured at 37 °C with an agitation of ~500 r.p.m. During the culturing of up to 12 h, the optical density (OD_595_) of each well were recorded by using a plate reader (Thermo multiskan FC) at a one-hour interval. The mean value and standard deviation of the recorded OD_595_ data (each repeated three times) were processed with the Python language and plotted with the Matplotlib software package (**Fig. 7**).

## Acknowledgements

We thank Professor Peter Schultz (The Scripps Research Institute, USA) for providing us with the plasmids that carry the genes encoding the orthogonal tRNA and amino acyl-tRNA synthetase for Bpa incorporation. We thank Keio Collections for providing us the BW25113 *E. coli* strain.

This work was supported by funds from the National Natural Science Foundation of China (No. 31971189, 31670775 and 31470766 to ZYC; No. 31770830 and No. 31570778 to X.F), the National Basic Research Program of China (No. 2012CB917300 to ZYC), and the Qidong-SLS Innovation Fund (to ZYC).

