## Supplemental Figure S1, Table for "High-throughput living-cell protein crosslinking analysis uncovers the physiological relevance of forming the “inserted” state of the ATP synthase ε-subunit"

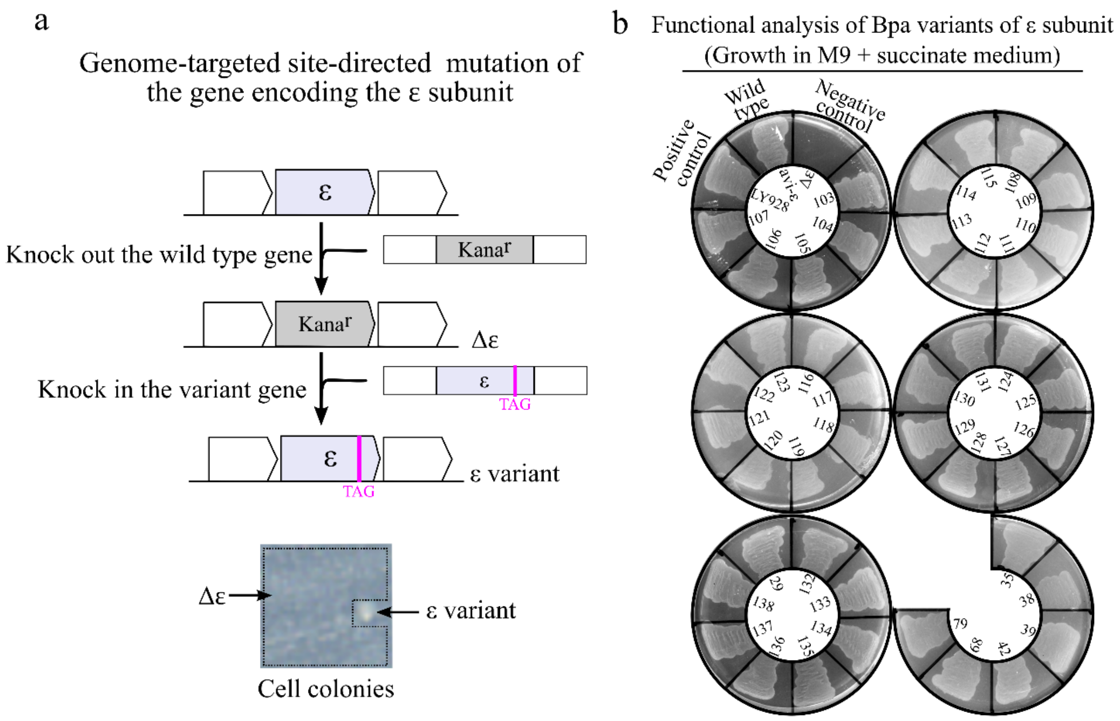


**Figure S1. Generation of the variants of the ε-subunit based on the genome-targeted site-directed mutagenesis technique and the phenotype analysis of the cells producing different variants of the ε-subunit.** (**a**) The site-directed mutagenesis was performed via a procedure involving two rounds of homologous recombination (top) and cells having their ε-subunit deleted ( Δ ε) and a mutated but functional ε-subunit (as the Bpa variants) will respectively have a defective (thus forming tiny-sized colonies) and normal growth (forming normal-sized colonies) in LB rich medium (bottom). (**b**) All the Bpa variants of the ε-subunit allowed the cells to grow in the succinate-containing M9 minimal medium, indicating the formation of a functional ATP synthase. The Δ ε cells were analyzed here as a negative control, while the LY928 and wild type cells (LY928 Avi-ε) were analyzed as positive controls.

**Table S1**

**Genotypes of all the *E. coli* strains used in this study.**

| Strains | Genotypes[a] | Sources |
|---|---|---|
| LY928 | BW25113 insH11::aminoacyl-tRNA synthetase of Bpa- $tRNA^{Bpa}$ | Laboratory storage |
| Δε | LY928 ε::$Kana^r$ | Recombineering |
| WT (aka. Avi-ε) | LY928 Avi-ε | Recombineering |

| | | |
|---|---|---|
| 104 | WT Avi-ε-104Bpa | Recombineering |
| 105 | WT Avi-ε-105Bpa | Recombineering |
| 106 | WT Avi-ε-106Bpa | Recombineering |
| 107 | WT Avi-ε-107Bpa | Recombineering |
| 108 | WT Avi-ε-108Bpa | Recombineering |
| 109 | WT Avi-ε 109Bpa | Recombineering |
| 110 | WT Avi-ε 110Bpa | Recombineering |
| 111 | WT Avi-ε 111Bpa | Recombineering |
| 112 | WT Avi-ε 112Bpa | Recombineering |
| 113 | WT Avi-ε 113Bpa | Recombineering |
| 114 | WT Avi-ε 114Bpa | Recombineering |
| 115 | WT Avi-ε 115Bpa | Recombineering |
| 116 | WT Avi-ε 116Bpa | Recombineering |
| 117 | WT Avi-ε 117Bpa | Recombineering |
| 118 | WT Avi-ε 118Bpa | Recombineering |
| 119 | WT Avi-ε 119Bpa | Recombineering |
| 120 | WT Avi-ε 120Bpa | Recombineering |
| 121 | WT Avi-ε 121Bpa | Recombineering |
| 122 | WT Avi-ε 122Bpa | Recombineering |
| 123 | WT Avi-ε 123Bpa | Recombineering |
| 124 | WT Avi-ε 124Bpa | Recombineering |
| 125 | WT Avi-ε 125Bpa | Recombineering |
| 126 | WT Avi-ε 126Bpa | Recombineering |
| 127 | WT Avi-ε 127Bpa | Recombineering |
| 128 | WT Avi-ε 128Bpa | Recombineering |
| 129 | WT Avi-ε 129Bpa | Recombineering |
| 130 | WT Avi-ε 130Bpa | Recombineering |
| 131 | WT Avi-ε 131Bpa | Recombineering |

| | | |
|---|---|---|
| 132 | WT Avi-ε 132Bpa | Recombineering |
| 133 | WT Avi-ε 133Bpa | Recombineering |
| 134 | WT Avi-ε 134Bpa | Recombineering |
| 135 | WT Avi-ε 135Bpa | Recombineering |
| 136 | WT Avi-ε 136Bpa | Recombineering |
| 137 | WT Avi-ε 137Bpa | Recombineering |
| 138 | WT Avi-ε 138Bpa | Recombineering |
| 121Bpa,I125L | WT Avi-ε 121Bpa,I125L | Recombineering |
| 121Bpa,I125V | WT Avi-ε 121Bpa,I125V | Recombineering |
| 121Bpa,I125F | WT Avi-ε 121Bpa,I125F | Recombineering |
| 121Bpa,I125E | WT Avi-ε 121Bpa,I125E | Recombineering |
| 121Bpa,I125K | WT Avi-ε 121Bpa,I125K | Recombineering |
| 121Bpa,I125G | WT Avi-ε 121Bpa,I125G | Recombineering |
| 121Bpa,I125A | WT Avi-ε 121Bpa,I125A | Recombineering |
| 121Bpa,I125N | WT Avi-ε 121Bpa,I125N | Recombineering |
| 121Bpa,I125D | WT Avi-ε 121Bpa,I125D | Recombineering |
| 121Bpa,I125T | WT Avi-ε 121Bpa,I125T | Recombineering |
| 121Bpa,I125S | WT Avi-ε 121Bpa,I125S | Recombineering |
| 121Bpa,I125Q | WT Avi-ε 121Bpa,I125Q | Recombineering |
| 121Bpa,I125M | WT Avi-ε 121Bpa,I125M | Recombineering |
| 121Bpa,I125Y | WT Avi-ε 121Bpa,I125Y | Recombineering |
| 121Bpa,I125W | WT Avi-ε 121Bpa,I125W | Recombineering |
| 121Bpa,I125H | WT Avi-ε 121Bpa,I125H | Recombineering |
| 121Bpa,I125C | WT Avi-ε 121Bpa,I125C | Recombineering |
| 121Bpa,I125R | WT Avi-ε 121Bpa,I125R | Recombineering |
| 121Bpa,I125P | WT Avi-ε 121Bpa,I125P | Recombineering |
| I125K | WT Avi-ε I125K | Recombineering |

[a]The codon “TAG” would be recognized and translated to the Bpa unnatural amino acid residue in LY928 strain and all the strains derived from it.
